# Single-chromosome aneuploidy commonly functions as a tumor suppressor

**DOI:** 10.1101/040162

**Authors:** Jason M. Sheltzer, Julie H. Ko, Nicole C. Habibe Burgos, Erica S. Chung, Colleen M. Meehl, Verena Passerini, Zuzana Storchova, Angelika Amon

**Affiliations:** David H. Koch Institute for Integrative Cancer Research and Howard Hughes Medical Institute, Massachusetts Institute of Technology, Cambridge, MA 02139.; Albert Einstein College of Medicine, Bronx, NY 10461.; Group Maintenance of Genome Stability, Max Planck Institute of Biochemistry, Martinsried, Germany; Cold Spring Harbor Laboratory, Cold Spring Harbor, NY 11724.

## Abstract

Whole-chromosome aneuploidy is a hallmark of human malignancies. The prevalence of chromosome segregation errors in cancer – first noted more than 100 years ago – has led to the widespread belief that aneuploidy plays a crucial role in tumor development. Here, we set out to test this hypothesis. We transduced congenic euploid and trisomic fibroblasts with 14 different oncogenes or oncogene combinations, thereby creating genetically-matched cancer cell lines that differ only in karyotype. Surprisingly, nearly all aneuploid cell lines divided slowly *in vitro*, formed few colonies in soft agar, and grew poorly as xenografts, relative to matched euploid lines. Similar results were obtained when comparing a near-diploid human colorectal cancer cell line with derivatives of that line that harbored extra chromosomes. Only a few aneuploid lines grew at close to wild-type levels, and no aneuploid line exhibited greater tumorigenic capabilities than its euploid counterpart. These results demonstrate that rather than promoting tumorigenesis, aneuploidy, particularly single chromosome gains, can very often function as a tumor suppressor. Moreover, our results suggest one potential way that cancers can overcome the tumor suppressive effects of aneuploidy: rapidly-growing aneuploid cell lines that had evolved *in vitro* or *in vivo* demonstrated recurrent karyotype changes that were absent from their euploid counterparts. Thus, the genome-destabilizing effects of single-chromosome aneuploidy may facilitate the development of balanced, high-complexity karyotypes that are frequently found in advanced malignancies.

## Introduction

In the early 20^th^ century, Theodor Boveri proposed that abnormal karyotypes altered the equilibrium between pro- and anti-proliferative cellular signals, and were therefore capable of transforming primary cells into cancer cells (Boveri, 2008). “Boveri’s hypothesis” was one of the first genetic explanations of cancer development, and it helped motivate a century of research into the origins and consequences of chromosome segregation errors. Since Boveri’s time, it has been established that approximately 90% of solid tumors and 75% of hematopoietic cancers display whole-chromosome aneuploidy (Weaver and Cleveland, 2006). However, the precise relationship between aneuploidy and tumorigenesis remains unclear.

A preponderance of current evidence supports Boveri’s hypothesis (Gordon et al., 2012; Holland and Cleveland, 2009). First, individuals with Down syndrome (trisomy 21) frequently develop pediatric leukemia, suggesting a clear link between the gain of chromosome 21 and leukemogenesis (Seewald et al., 2012). Secondly, many human cancers exhibit recurrent aneuploidies (Ozery-Flato et al., 2011; Zack et al., 2013), and computational modeling has suggested that these patterns of chromosomal alterations reflect an evolutionary process in which cancer cells increase the copy number of loci encoding oncogenes and decrease the copy number of loci encoding tumor suppressors (Davoli et al., 2013). Finally, genetically-engineered mice that harbor alleles which cause chromosomal instability (CIN) typically develop tumors at accelerated rates (Jeganathan et al., 2007; Li et al., 2009; Michel et al., 2001; Park et al., 2013; Sotillo et al., 2007, 2010), particularly when combined with mutations in the p53 tumor suppressor (Li et al., 2010). Low levels of CIN have been reported to be particularly tumorigenic (Silk et al., 2013). Nonetheless, several observations suggest that the relationship between aneuploidy and cancer may be more complex than previously believed. While individuals with Down syndrome are at an increased risk of developing leukemia and germ cell tumors, they are at a significantly decreased risk of developing many other common solid tumors (Nižetić and Groet, 2012). Trisomies of regions orthologous to human chromosome 21 in the mouse have also been found to suppress tumor development (Baek et al., 2009; Reynolds et al., 2010; Sussan et al., 2008). Moreover, though mouse models of CIN are generally tumor-prone, in certain organs or when combined with certain oncogenic mutations, CIN mice exhibit reduced tumor burdens (Silk et al., 2013; Weaver et al., 2007). Thus, aneuploidy may have tumor-protective as well as tumor-promoting effects, which could differ depending on the genetic and environmental milieu.

In order to further our understanding of the effects of aneuploidy on cell and organismal physiology, systems have been developed to generate primary cells with a range of aneuploid karyotypes (Pavelka et al., 2010; Stingele et al., 2012; Torres et al., 2007; Williams et al., 2008). These cells have been constructed without CIN-promoting mutations, thereby allowing the study of aneuploidy absent other genetic perturbations. This research has demonstrated the existence of a set of phenotypes that are shared among many different aneuploid cells and are largely independent of the specific chromosomal alteration: aneuploid cells display reduced fitness (Stingele et al., 2012; Torres et al., 2007; Williams et al., 2008), are deficient at maintaining proteostasis (Donnelly et al., 2014; Oromendia et al., 2012; Tang et al., 2011), and exhibit a specific set of gene expression changes that include the down-regulation of cell-cycle transcripts and the up-regulation of a stress-response program (Dürrbaum et al., 2014; Sheltzer, 2013; Sheltzer et al., 2012). A crucial question, however, is in what way(s) the cellular changes induced by aneuploidy affect (and possibly drive) tumorigenesis. Aneuploid cells may be poised to undergo transformation due to their increased dosage of oncogenes and decreased dosage of tumor suppressors (Davoli et al., 2013), the inherent instability of aneuploid genomes (Duesberg et al., 1999), or due to a general misregulation of cell metabolism and other biological processes (Rasnick and Duesberg, 1999). To test these ideas, we compared the tumorigenicity of a series of genetically-matched euploid and aneuploid cells. Surprisingly, we found that nearly every aneuploid cell line that we examined displayed reduced tumor-forming potential relative to control euploid cell lines. These results necessitate a significant revision of our understanding of the relationship between aneuploidy and cancer.

## Results

### Single-chromosome aneuploidy is insufficient to induce neoplastic phenotypes

We took advantage of naturally-occurring Robertsonian translocations to generate mouse embryonic fibroblasts (MEFs) trisomic for chromosome 1, 13, 16, or 19, as well as sibling-matched euploid controls (Williams et al., 2008). While advanced malignancies frequently harbor complex karyotypes that include multiple chromosome gains and/or losses, early-stage cancers typically exhibit one or a few arm-length or whole-chromosome aneuploidies (Balaban et al., 1986; Di Capua Sacoto et al., 2011; Lai et al., 2007; Magnani et al., 1994; El-Rifai et al., 2000). Thus, these trisomies likely recapitulate the karyotypic state of pre-malignant or early-stage cancer lesions, and their study can shed light on the role of aneuploidy in cancer development and evolution.

Read depth analysis from low-pass whole genome sequencing of each MEF line demonstrated that these cells harbored complete trisomies without other chromosomal alterations and that the extra chromosomes were present clonally within the cell population (Figure 1A). Various oncogenes are encoded on these chromosomes, including BCL2 (mChr1), FGFR4 (mChr13), Jak2 (mChr19), and many others (Table S1). Gain of these oncogenes, or some other consequence of aneuploidy, could drive malignant growth or otherwise generate cancer-like phenotypes in primary cells. We therefore set out to discover whether single-chromosome aneuploidy in MEFs was sufficient to induce neoplastic or pre-neoplastic behavior in untransformed cells.

**Figure 1.**
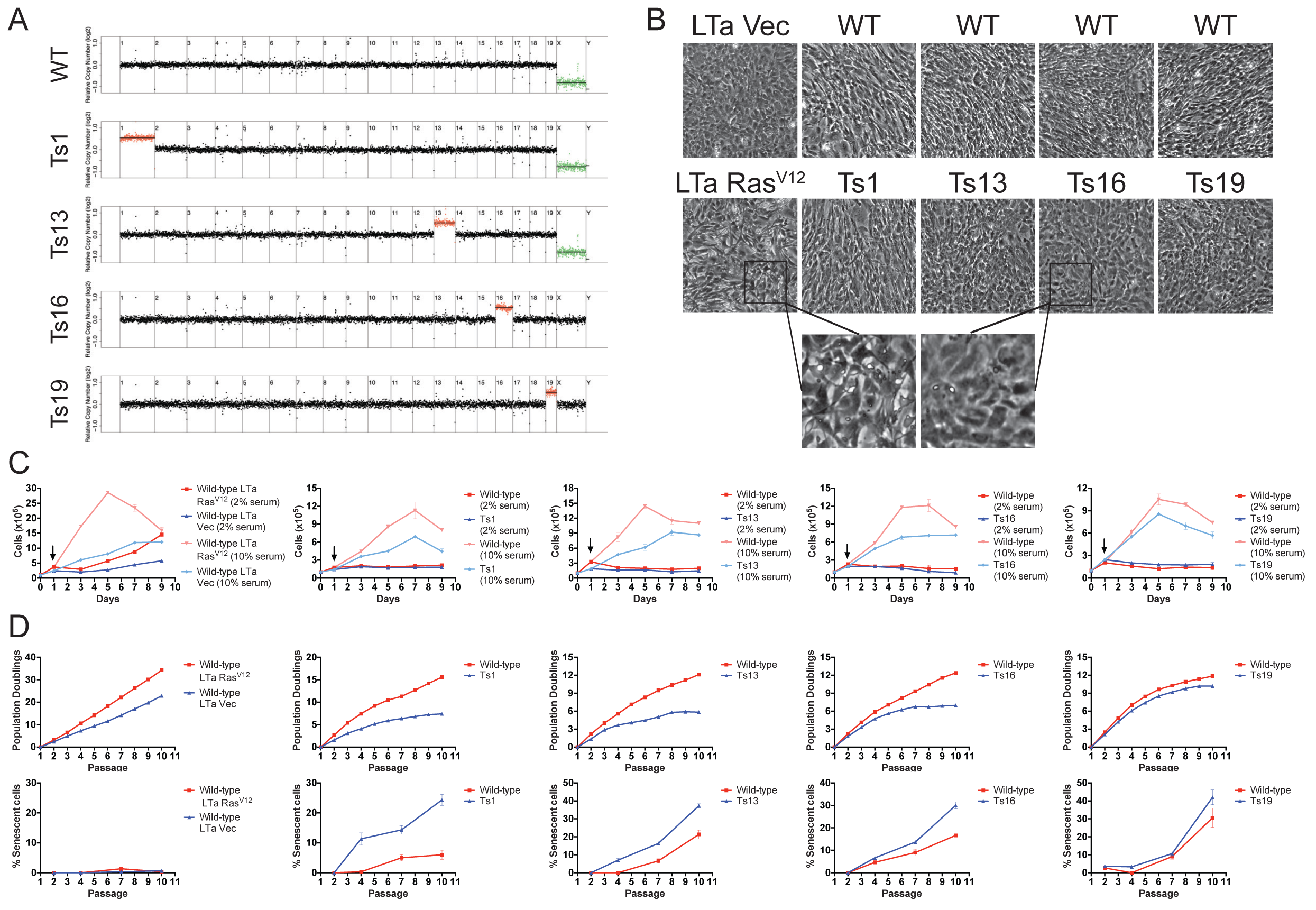
Single-chromosome aneuploidy is insufficient to induce neoplastic phenotypes. (A) MEF lines that were euploid or trisomic for chromosomes 1, 13, 16, or 19 were subjected to low-pass whole genome sequencing. Normalized read depths across 500kb bins are displayed. Note that only one euploid cell line is shown, although each trisomic MEF line had a separate euploid line that was derived from a euploid littermate. (B) Photomicrographs of monolayers of the indicated cell lines. LTa+Ras^V12^ MEFs, but not trisomic MEFs, lose contact inhibition when grown to confluence. (C) Growth curves of the indicated cell lines are displayed. Cells were first plated in normal (10% serum) medium, then 24 hours after plating the cells were re-fed or switched to reduced (2% serum) medium (indicated by an arrow). LTa+Ras^V12^ MEFs, but not trisomic MEFs, continue to divide in low serum media. (D) The indicated cells were passaged, counted, and plated in triplicate every third day for 10 passages (top row). On passages 2, 4, 7, and 10, β-galactosidase levels were measured (bottom row). LTa-transduced MEFs exhibit negligible levels of senescence, but trisomic cell lines senesce at an early passage.

As a positive control for cancer-like growth, we generated MEF lines that had been stably-transduced with the Large T antigen (LTa), which inhibits the Rb and p53 tumor suppressor pathways (Ahuja et al., 2005), and with either an empty vector or an activated allele of H-Ras (Ras^V12^). As expected, the LTa+Ras^V12^ MEFs exhibited several neoplastic phenotypes: the cells were not contact-inhibited, and instead piled on top of each other when grown to confluence (Figure 1B), they formed colonies from single cells when plated at low density (Figure S1), they grew independently of pro-proliferative signals, as evidenced by their increase in cell number when plated in low-serum medium (Figure 1C), and they grew robustly without senescing over 10 passages in culture (Figure 1D). The LTa+vector MEFs displayed an intermediate, pre-neoplastic phenotype: the cells maintained contact inhibition and grew very poorly following serum withdrawal, but were mildly clonogenic and doubled without noticeable senescence over 10 passages in culture. In contrast, both euploid and trisomic MEFs failed to display any cancer-like phenotypes: they exhibited appropriate contact inhibition, failed to proliferate in low-serum medium, were non-clonogenic, and senesced after 7 to 10 passages in culture (Figure 1 and S1). Interestingly, trisomies 1, 13, and 16 senesced at earlier passages and to a significantly higher degree than their matched euploid lines, as judged by β-galactosidase staining (Figure 1D). The proliferation defect and increased senescence of the aneuploid cell lines was approximately proportional to the degree of aneuploidy: cells trisomic for mChr1 (the largest mouse autosome) displayed the most severe phenotypes, while trisomy of mChr19 (the smallest mouse autosome) induced more subtle effects. We conclude that single-chromosome aneuploidy is insufficient to generate neoplastic phenotypes, and many aneuploidies in fact induce a premature growth arrest.

### Oncogene-transduced trisomic cells exhibit reduced proliferation and fitness relative to oncogene-transduced euploid cells

As aneuploidy alone was insufficient to generate cancer-like phenotypes, we next set out to determine whether aneuploidy could have synergistic, growth-promoting interactions with oncogenic mutations. In particular, loss of the p53 tumor suppressor has been linked with heightened proliferation of aneuploid cells (Thompson and Compton, 2010). However, these studies did not examine the effects of loss of p53 in euploid cells, leaving unresolved the question of whether or not the proliferative benefits conferred by the loss of p53 are specific to (or relatively greater in) aneuploid cells. To address this question, and to screen for synergistic interactions between aneuploidy and common oncogenic mutations, we stably transduced trisomic and euploid MEFs with plasmids encoding various oncogenes or with matched empty vectors. Trisomies 1, 13, 16, and 19, as well as matched euploid cell lines, were transduced with a dominant negative allele of TP53 [p53dd (Shaulian et al., 1992)], the E1a oncogene [which inhibits the Rb tumor suppressor, among several other cellular pathways (Gallimore and Turnell, 2001)], the Large T oncogene, or with the MYC oncogene. Low-pass whole genome sequencing at early passage following transduction showed that the cell lines maintained their initial karyotypes, confirming that we had constructed oncogene-expressing cell lines with single, defined chromosomal copy number alterations (Figure S2A). Following retroviral transduction and selection, the behavior of euploid and trisomic cell lines was tested in several assays that serve as *in vitro* and *in vivo* proxies for tumorigenic capacity.

Oncogene-transduced euploid and trisomic MEFs were counted and passaged in a modified 3T3 protocol 10 times over the course of 30 days following selection. Empty vector-transduced trisomic MEFs generally underwent fewer population doublings than matched euploid lines (Figure 2A, compare dark red and dark blue lines). Transduction with oncogenes significantly enhanced the growth of trisomic cells, and the resultant lines doubled more frequently over the course of the experiment than the vector-matched controls. However, these oncogenes also enhanced the growth of euploid cell lines. In the majority of cases, the oncogene-transduced euploid cell line underwent more population doublings than the corresponding oncogene-transduced aneuploid line, and in only a few instances did we observe equivalent proliferation between euploid and trisomic MEFs (Figure 2A, compare light red and light blue lines). For instance, over 10 passages in culture, p53dd-transduced trisomy 13 cells doubled about 9 times, while a p53dd-transduced euploid line doubled 15 times. Cells trisomic for chromosome 1 (the largest mouse autosome) generally grew the slowest relative to wild-type, while cells trisomic for chromosome 19 (the smallest mouse autosome) tended to proliferate at close to wild-type levels. Overall, expression of p53dd or MYC failed to suppress the growth differential between aneuploid and euploid cell lines, while the expression of E1a or LTa had more potent effects. Transduction with Large T, in particular, demonstrated the most significant suppression of the aneuploidy-induced proliferation defect, although LTa-expressing cells trisomic for chromosomes 1 and 16 still divided significantly more slowly than LTa-expressing euploid lines.

**Figure 2.**
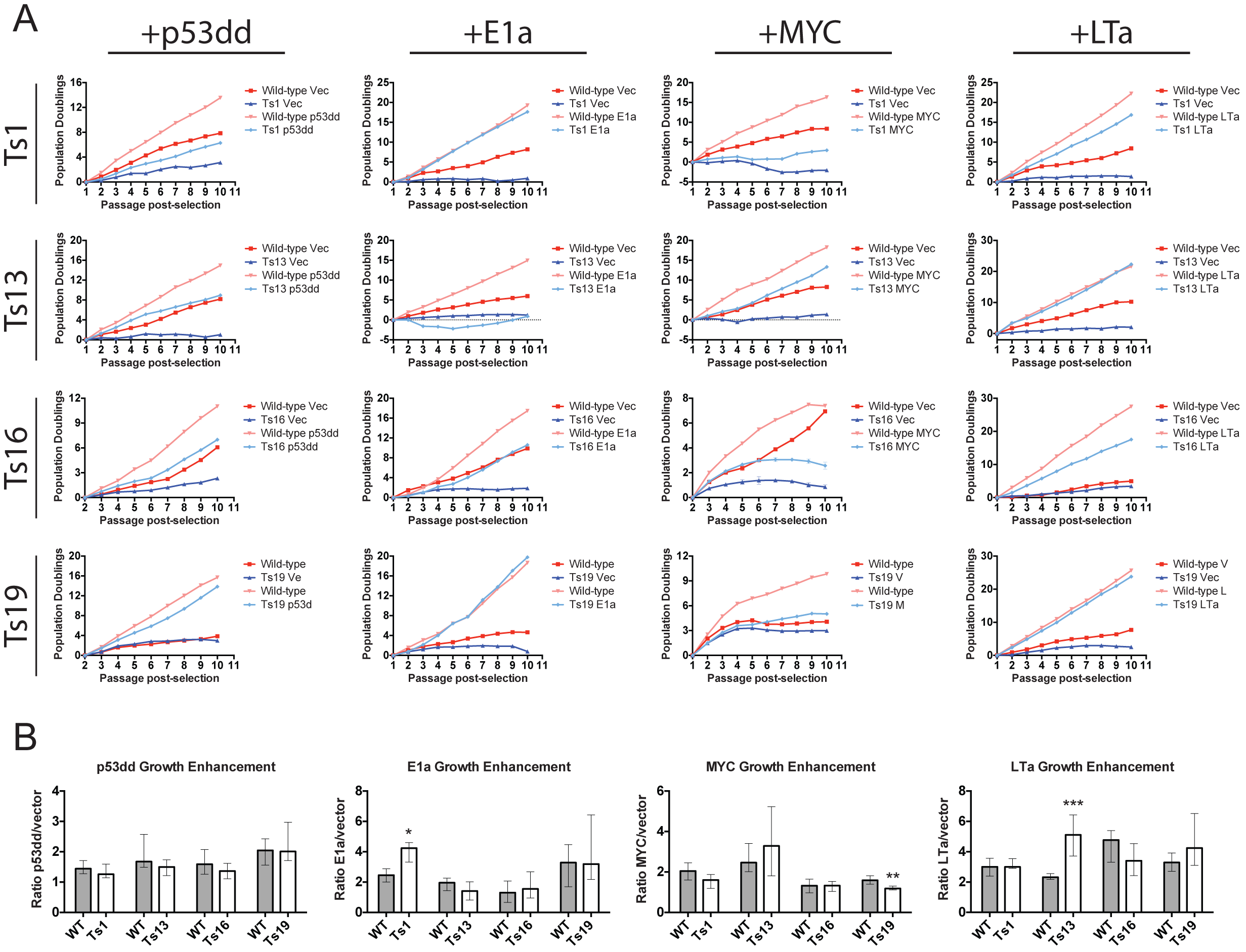
Aneuploidy impedes the proliferation of oncogene-transduced cell lines. (A) Euploid and trisomic cell lines were stably transduced with plasmids harboring the indicated oncogene or a matched empty vector. Following selection, the cell lines were passaged every third day for up to 10 passages, and the cumulative population doublings over the course of each experiment are displayed. Note that the panel displaying Ts19+LTa is reproduced in Figure 7. (B) The number of cells recovered from oncogene-transduced MEFs was divided by the number of cells recovered from vector-transduced MEFs at every passage. Bar graphs display the median ratios and the interquartile ranges. * p<.05; ** p<.005 (Wilcoxon rank-sum test).

We next tested several additional oncogenes in a subset of trisomies: euploid cell lines transduced with a stabilized allele of MYC (MYC^T58A^) outgrew matched cell lines trisomic for chromosomes 13 or 16, while expression of an activated allele of BRAF (BRAF^V600E^) or an allele of CDK4 resistant to p16 inhibition (CDK4^R24C^) resulted in senescence in both euploid and trisomic cell lines (Figure S3).

As the experiments described above were conducted using initial populations of primary cells, we selected nine experiments to repeat on independently-derived cell lines (Figure S4). Replicate experiments displayed some line-to-line variability (e.g., compare Ts16+LTa), but recapitulated the major features of our initial results: oncogene-transduced trisomic cells grew less rapidly than euploid cells, and the expression of Large T provided the most significant rescue of trisomic cell growth. (Figure S4).

The prolonged culture period also allowed us to follow the dynamics of aneuploid cell populations over time: rapidly-dividing subpopulations could potentially arise during 30 days in culture that would enhance the apparent proliferative capacity of the aneuploid MEFs. Interestingly, in our replicate analysis on independently-derived cell lines, we found that the proliferation rate of one Ts19+LTa cell line increased during serial passaging, while an independent Ts19+LTa line grew at approximately the same rate over the course of the experiment (compare Figure 2A and Figure S4). The reasons for this divergent behavior are explored below. In general, few trisomic cell lines doubled more rapidly over successive passages, while many trisomic cell lines grew more slowly over time instead. Consistent with their poor proliferation at high passage, p53‐ and MYC-transduced trisomic lines displayed elevated levels of senescence-associated β-galactosidase staining relative to euploid controls (Figure S5A). However, expression of E1a or LTa effectively blocked senescence in all cells, though a fitness differential between euploid and trisomic populations was still evident, suggesting that other factors contribute to the growth delay in aneuploid cells. Across all of our experiments, we did not observe any instances in which transduction with an oncogene generated a trisomic cell line that consistently outgrew its matched euploid counterpart across multiple independent lines. We conclude that euploid lines harboring common oncogenic mutations generally proliferate more rapidly than trisomic lines harboring the same genetic alterations.

Though proliferative differences between aneuploid and euploid lines persisted following oncogene transduction, it remained possible that the oncogenes provided a relatively greater growth advantage to aneuploid cells than to euploid cells. To test this, we quantified the benefit conferred by each oncogene by comparing the number of cells recovered at every passage from oncogene-transduced and vector-transduced cell lines (Figure 2B). For most oncogene-trisomy combinations, the fold change in growth enhancement was equivalent between euploid and aneuploid lines. For instance, transduction with dominant-negative p53 resulted in an approximately 1.6 to 1.8-fold increase in doublings per passage, relative to the vector-transduced controls, in all euploid and trisomic cell lines. Thus, the abrogation of p53 signaling does not specifically enhance the growth of these trisomic cells. In total, across 16 oncogene-aneuploidy combinations, oncogene expression was found to confer a similar proliferative benefit to euploid and trisomic cells in 13 combinations, while in only 2 conditions (Ts1+E1a and Ts13+LTa) oncogene expression was found to have a stronger effect on the trisomic MEFs than on the euploid MEFs. These results suggest that oncogene-aneuploidy synergy is rarely observed, and may be specific to certain chromosomes or oncogenes.

As an additional test of the proliferative capabilities of the oncogene-transduced lines, we assessed the focus formation ability of cells that had been transduced with E1a or LTa (MYC and p53dd-transduced lines remained non-clonogenic). We found that transduced euploid lines exhibited uniformly superior colony-forming ability relative to the trisomic lines, even in instances when the euploid and aneuploid lines demonstrated approximately equal doubling times in culture (Figure S7). For instance, while Ts19+LTa and WT+LTa MEFs grew at nearly the same rate, the WT MEFs formed about 6-fold more colonies when plated as single cells than the trisomic line did. The differences between the colony formation and population doubling assays likely reflect the fact that forming a colony is a relatively greater challenge to a cell than doubling in a monolayer, and this challenge exacerbates the fitness differential between euploid and trisomic cells. In summary, our data indicate that single-chromosome gains cause a pervasive fitness defect, even in oncogene-transduced populations.

### Ras^V12^-transformed aneuploid cells exhibit reduced tumorigenicity

Singly-transduced MEFs are not fully-transformed; complete transformation of rodent cells requires transduction with two oncogenes (Land et al., 1983). We therefore took euploid and trisomic p53dd‐ or LTa-transduced cell lines and then stably transduced them with a vector harboring oncogenic H-Ras^V12^ or a control empty vector. Whole genome sequencing revealed that 14 out of 14 tested cell lines maintained their initial karyotype following two rounds of retroviral transduction and selection, demonstrating that we had successfully generated identically-transformed cell lines with single-chromosome differences (Figure S2B).

Ras^V12^-transduced cell lines doubled significantly faster than vector-transduced lines over 10 passages in culture, and Ras^V12^ narrowed or in some cases abolished the proliferative difference between euploid and trisomic MEFs (Figure 3A). For instance, while p53dd+vector-transduced WT and Ts13 MEFs doubled 20 and 6 times, respectively, p53dd+Ras^V12^-transduced WT and Ts13 MEFs doubled 32 and 30 times, respectively. However, no Ras^V12^-transduced trisomic line grew faster than its euploid counterpart, and in subsequent assays described below, a fitness benefit conferred by euploidy was clearly detected. Ras^V12^-transduced trisomic cell lines were also found to display elevated the levels of senescence-associated β-galactosidase, even when co-expressed with LTa (Figure S5A). Additionally, we examined the effects of Ras^V12^ expression in E1a‐ or MYC-transduced Ts16 cell lines (Figure S8), and we tested the expression of BRAF^V600E^, PIK3CA^H1047R^, or MYC in lieu of Ras^V12^ as a driver oncogene (Figure S8). The latter oncogenes typically had little effect in this assay compared to experiments using Ras^V12^ that were performed in parallel, and no other oncogene exhibited a strong suppression of the aneuploidy-induced growth delay.

**Figure 3.**
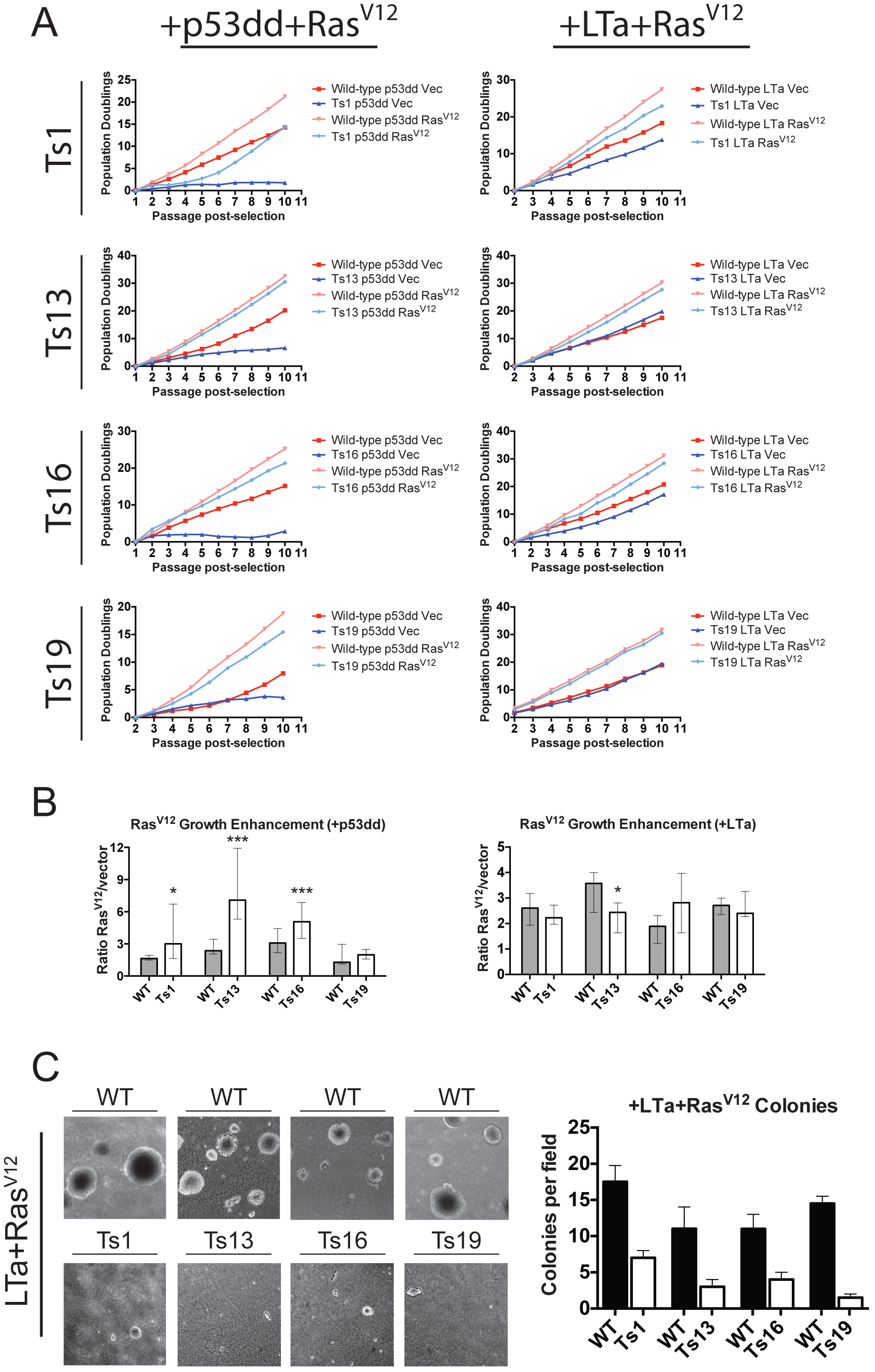
Effects of Ras^V12^ expression on immortalized euploid and trisomic MEFs. (A) Euploid and trisomic cell lines were first stably transduced with p53dd or with LTa, and then transduced a second time with plasmids harboring Ras^V12^ or a matched empty vector. The cell lines were passaged, counted, and plated in triplicate up to 10 passages following the second round of selection. Note that the panel displaying Ts1+p53dd+Ras^V12^ is reproduced in Figure S12, the panel displaying Ts13+LTa+Ras^V12^ is reproduced in part in Figure S8, the panel displaying Ts16+p53dd+Ras^V12^ is reproduced in Figure S9, and the panel displaying Ts19+LTa+RasV12 is reproduced in part in Figure S8 and Figure S11. (B) The number of cells recovered from Ras^V12^-transduced MEFs was divided by the number of cells recovered from vector-transduced MEFs at every passage. Bar graphs display the median ratios and the interquartile ranges. * p<.05; *** p<.0005 (Wilcoxon rank-sum test). (C) 20,000 cells of the indicated cell lines were plated in soft agar and then grown for 20 days. For each comparison, the euploid MEFs formed more colonies than the trisomic MEFs (p<.01, Student’s t test).

In order to assess the reproducibility of our results, we performed several replicate experiments on independently derived and immortalized cell lines (Figure S8). One line of Ts19+LTa+PIK3CA^H1047R^ grew slightly better than its euploid control, though an independent line of Ts19+LTa+PIK3CA^H1047R^ did not exhibit this phenotype (Figure S8, and see Figure S11 below). In other replicate experiments, we also observed some variability in the degree of growth rescue caused by oncogene expression. For instance, in an initial experiment, p53dd+Ras^V12^ transduction resulted in an incomplete suppression of the proliferative difference between WT and Ts16 cells, while in a replicate experiment, strong suppression was induced by this oncogene combination. To address the origins and scope of this variability, we repeated this experiment a total of 6 times using 5 independent cell lines (Figure S9). When the same primary cell line was transduced with p53dd and Ras^V12^, both sets of transformed lines exhibited comparable proliferation dynamics. When four other Ts16 cell lines were used, each transformed line grew at a moderately different rate, though no trisomic line was observed to divide more rapidly than its euploid counterpart (Figure S9). We conclude that variability within this assay results from inherent differences between primary cell lines, rather than from stochasticity in the experimental protocol. Moreover, while the growth penalty induced by aneuploidy varies among independently-derived cell lines, across multiple replicate experiments, no oncogene cocktail tested resulted in reproducibly superior growth in a trisomic line compared to its euploid counterpart.

In order to determine the relative benefit conferred by Ras^V12^ expression to euploid and aneuploid MEFs, we compared the number of cells recovered at each passage from Ras^V12^-transduced and vector-transduced cell lines. Ras^V12^ expression conferred a similar fold-increase in proliferative capacity in euploid and trisomic MYC-, E1A-, and three LTa-transduced cell lines, while Ras^V12^ expression had a proportionately greater effect on WT+LTa lines than on Ts13+LTa lines (Figure 3B and Figure S6). Interestingly, in a p53dd background, Ras^V12^ had a greater effect on trisomies 1, 13, and 16 than it did on the euploid cell lines. This is likely due at least in part to increased senescence of the p53dd+vector doubly-transduced trisomic MEFs (Figure S5A).

As the Ras^V12^-transduced trisomic lines displayed equivalent or nearly-equivalent proliferative capacity as the euploid lines, and as Ras^V12^ expression showed some evidence of synergy with p53 inactivation in trisomic MEFs, we hypothesized that Ras^V12^ suppresses the fitness differential between aneuploid and euploid cells. However, this suppression was less evident in other assays for tumorigenicity. Ras^V12^-transduced euploid MEFs formed more colonies from single cells than trisomic lines did, even when the lines were observed to proliferate at the same rate in culture (Figure S10). Fully transformed cell lines are also competent to grow in soft agar, a phenotype that strongly correlates with *in vivo* tumorigenicity (Shin et al., 1975). We tested the ability of LTa+Ras^V12^-transduced cell lines to form colonies in soft agar, and in each experiment the euploid control lines exhibited higher colony-forming ability than the equivalently-transduced trisomic lines (Figure 3C).

As a final test of the tumorigenicity of transformed trisomic MEFs, we examined the ability of Ras^V12^-transduced euploid and trisomic cell lines to form tumors in xenograft assays. Euploid and trisomic p53dd+Ras^V12^ cell lines were injected contralaterally into the flanks of nude mice, and tumor volume measurements were obtained every third day. While aneuploid and euploid cell lines grew at similar rates *in vitro*, the euploid lines invariably formed significantly larger tumors *in vivo* (Figure 4).

**Figure 4.**
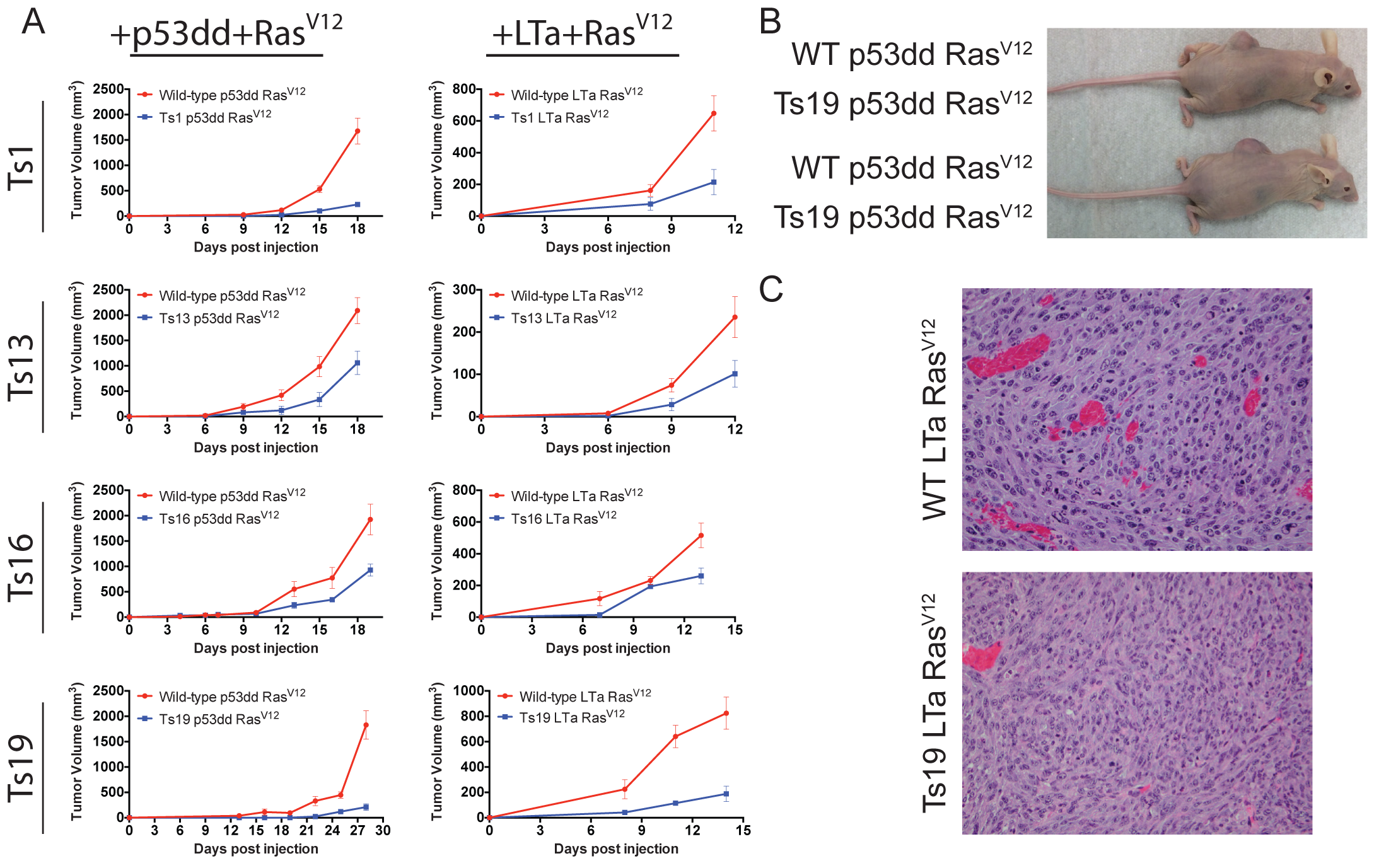
Trisomy hampers tumor growth in xenografts. (A) 10^6^ euploid or aneuploid cells transduced with either p53dd and Ras^V12^ or with LTa and Ras^V12^ were injected subcutaneously into the flanks of nude mice. Tumor volume was measured every three days. Note that mice injected with cells transduced with LTa+Ras^V12^ had to be euthanized prematurely due to cachexia. (B) Representative images of mice injected contralaterally with WT+p53dd+Ras^V12^ cells or Ts19+p53dd+Ras^V12^ cells. (C) Representative sections of xenograft tumors from WT+LTa+Ras^V12^ cells or Ts19+LTa+Ras^V12^ cells stained with H&E.

We also examined the tumorigenicity of trisomic cell lines that had been transduced with LTa and Ras^V12^. In preliminary experiments, a fraction of mice injected with these cell lines developed cachexia that resulted from metastatic disease. In order to more easily identify the cell line that the metastases were derived from, euploid and trisomic cell lines were injected into different mice during single experiments. Additionally, mice were euthanized at 11 to 15 days post-injection, as cachexia began to develop. Consistent with our previous results, euploid LTa+Ras^V12^ cell lines formed larger tumors than trisomic LTa+Ras^V12^ cell lines. Following euthanasia, necropsies were performed on 29 mice: 3 out of 14 mice injected with trisomic cells and 5 out of 15 mice injected with euploid cells exhibited evidence of gross metastases (p=.68, Fisher’s exact test). Metastatic lesions were commonly observed on the stomach, spleen, liver, and pancreas of these mice, with no apparent difference in organ colonization between mice that had been injected with transformed euploid or trisomic cell lines (data not shown). Histological analysis identified the primary tumors and metastatic lesions as poorly-differentiated fibrosarcomas, consistent with their embryonic fibroblast origins (Figure 4C). No gross differences in histology were apparent between euploid and trisomic tumors. In total, these results demonstrate that tumors derived from euploid and trisomic cells form histologically similar structures, but euploid cells typically outgrow genetically-identical trisomic cells in xenografts.

### Chromosome gains impede the tumorigenicity of human colorectal cancer cells

Our experiments in MEFs demonstrated that aneuploidy in primary cells commonly impedes transformation. As aneuploidy is a nearly universal occurrence in cancer, we hypothesized that the acquisition of aneuploidy in previously-transformed cells, rather than in primary cells, could have distinct, pro-tumorigenic consequences. To test this, we utilized a series of chromosomally-stable human colorectal cancer cells into which extra chromosomes had been introduced via microcell-mediated chromosome transfer (Donnelly et al., 2014; Stingele et al., 2012). We first characterized each cell line by whole-genome sequencing (Figure 5A). Read-depth analysis of the parental cell line HCT116 confirmed several previously-described segmental gains on chromosomes 8, 10, 16, and 17. These amplifications were also present in the derived cell lines, and are therefore unlikely to affect the results described below.

**Figure 5.**
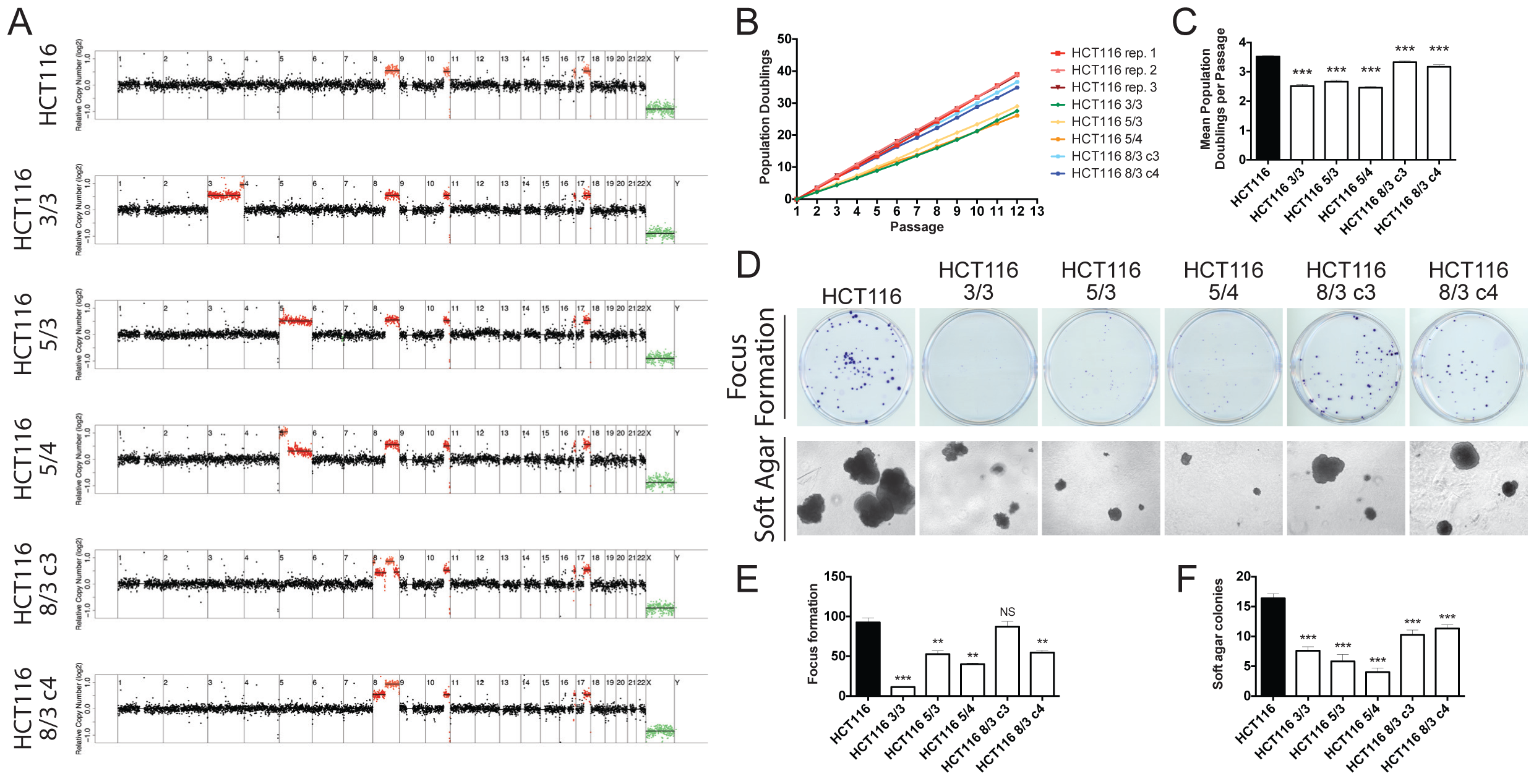
Aneuploidy impedes the growth of human colorectal cancer cell lines *in vitro*. (A) Normalized read depths from whole genome sequencing of the HCT116 human colorectal cancer cell line as well as HCT116 derivatives that harbored extra chromosomes. (B) Growth curves of colorectal cancer cell lines with different karyotypes. Cells were counted and passaged every third day. (C) Quantification of the mean population doublings per passage of multiple replicates of the experiment shown in (B). (D) 200 cells of the indicated lines were grown for 14 days prior to staining with crystal violet (top), or 2000 cells of the indicated lines were plated in soft agar and allowed to grow for 20 days before being imaged (bottom). (E) Quantification of focus formation assayed in (D). **, p<.005; ***, p<.0005 (Student’s t test). (F) Quantification of colony formation in soft agar in (D). **, p<.005; ***, p<.0005 (Student’s t test).

We compared the behavior of the parental HCT116 line to a cell line that was trisomic for chromosome 5 (HCT116 5/3), a cell line that had regions of trisomy and tetrasomy on chromosome 5 (HCT115 5/4), a cell line that had regions of trisomy and tetrasomy on chromosome 3 (HCT116 3/3), and two cell lines that had regions of trisomy and tetrasomy on chromosome 8 (HCT116 8/3 c4, which had gained a complete extra copy of chromosome 8, and HCT116 8/3 c3, which had gained a partial copy of chromosome 8). Oncogenes encoded on these chromosomes include β-catenin (hChr3), PIK3CA (hChr3), TERT (hChr5), MYC (hChr8), and several others (Table S2). We tested the HCT116 cell lines in similar assays as described above for the trisomic MEFs in order to compare the relative fitness and tumorigenicity of the parental and the derived colon cancer cell lines. During serial passaging, the near-diploid parental line divided the most rapidly, while the HCT116 3/3, 5/3, and 5/4 lines underwent on average one fewer doubling per passage (Figure 5B and 5C). HCT116 8/3 c3 and c4 divided at nearly the same rate as the wild-type line, although over the course of the experiment a small but significant growth delay was evident in these lines. Independent repetitions of this experiment produced identical results (Figure 5C). Parental HCT116 cells were also found to exhibit the highest rates of focus formation and colony growth in soft agar (Figure 5D-5F). HCT116 3/3, 5/3, and 5/4 displayed significant impairments in both assays. HCT116 8/3 c4 exhibited a small reduction in both assays, while HCT116 8/3 c3 grew moderately worse in soft agar but was able to form foci on plastic dishes at wild-type levels. We conclude that the introduction of aneuploidy into cancer cells commonly antagonizes growth *in vitro*, though in some instances it can be a nearly neutral event.

Consistent with our observations in MEFs, aneuploidy caused an increase in the levels of senescence-associated β-galactosidase expression in the human colorectal cancer cells. While the parental line and both hChr8 trisomies displayed minimal senescence, the level of staining was greatly increased in HCT116 3/3 and slightly increased in HCT116 5/3 and 5/4 cells (Figure S5B). The different levels of senescence between HCT116 3/3, 5/3, and 5/4 – all of which grow at similarly slow rates – further suggests that factors in addition to senescence contribute to the growth differential between the parental line and the derived aneuploid lines.

To determine how aneuploidy influenced tumorigenesis in these cell lines *in vivo*, we performed contralateral subcutaneous injections of either the parental HCT116 line or the aneuploid derivative lines into flanks of nude mice. The parental HCT116 cell line formed large tumors in all animals into which it had been injected (Figure 6). These tumors grew at a rapid rate, and each animal had to be euthanized 30 to 35 days after injection due to tumor burden. HCT116 3/3 and 5/4 formed small nodules at the site of injection that remained stable or increased in size only very slightly over the course of the experiment. Mice injected with HCT116 5/3 developed tumors that grew faster than either HCT116 3/3 or HCT116 5/4 but significantly less rapidly than the HCT116 tumors. Finally, consistent with our *in vitro* experiments, HCT116 8/3 c3 and c4 formed large tumors as xenografts, and there was no significant difference in tumor volume between these lines and the wild-type line. We conclude that gain of chromosomes in a cancer cell line can antagonize tumor formation.

**Figure 6.**
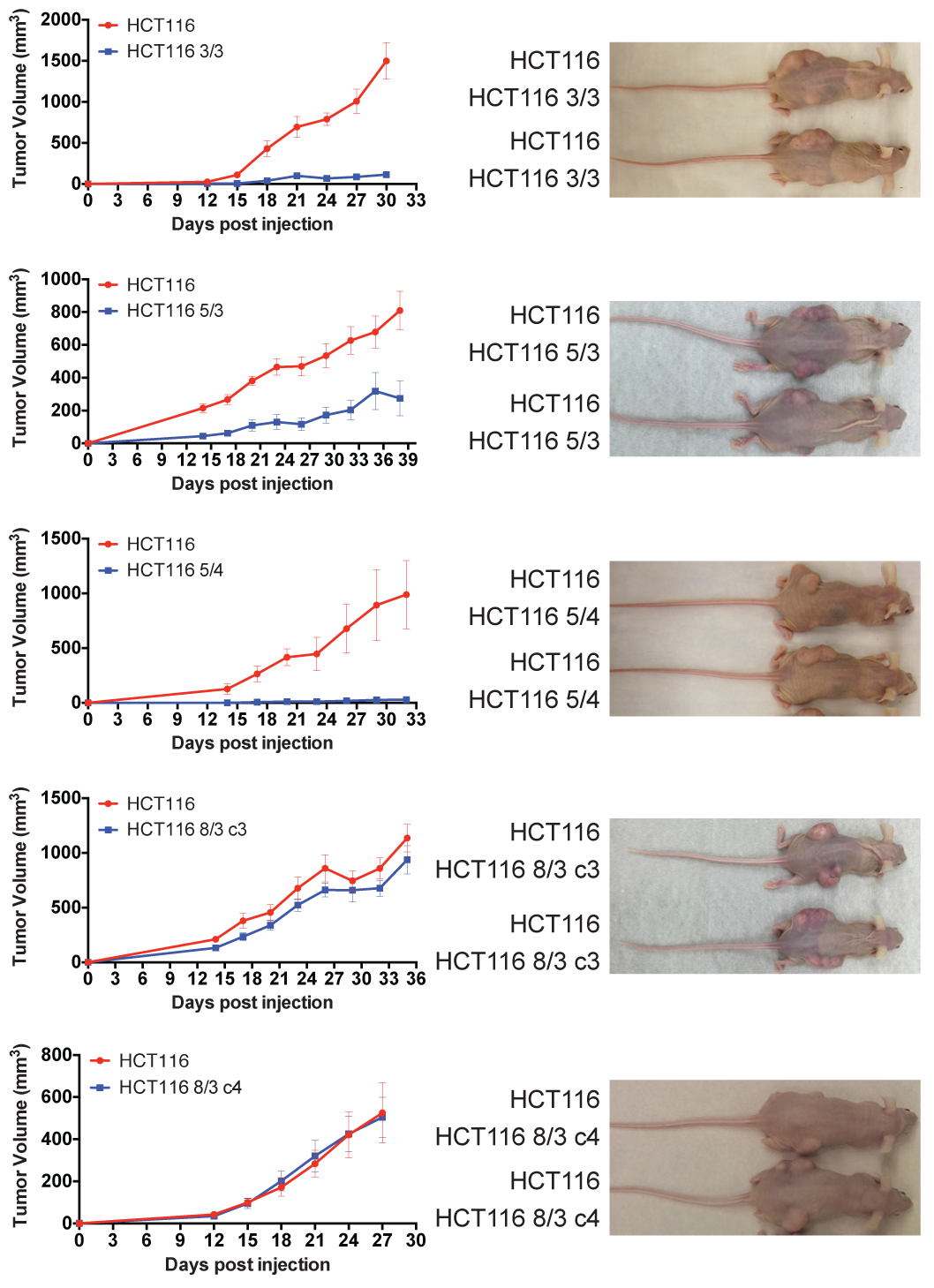
Aneuploidy impedes the growth of human colorectal cancer cell lines *in vivo*. 4×10^6^ HCT116 cells or HCT116 cells with additional chromosome(s) were injected subcutaneously into the flanks of 5-10 nude mice. Tumor growth was measured every third day.

### Improved growth in aneuploid cells is associated with further karyotypic alterations

The robust growth of aneuploid tumors suggests that cancer cells are able to adapt to the adverse effects of aneuploidy. Our results argue that one commonly-hypothesized aneuploidy-tolerating mechanism ‐ the activation of oncogenes and the inactivation of p53 and other tumor suppressors - is largely insufficient to equalize growth between euploid and aneuploid cells. We therefore sought to uncover other changes that could explain how cells are able to adapt to the aneuploid state.

In the oncogene-transduction experiments conducted in trisomic MEFs, we noted that one Ts19+LTa cell line initially proliferated at approximately the same rate as the euploid control line, but following several passages in culture, its proliferative rate increased (Figure 7A and Figure S4). An independent LTa-expressing Ts19 cell line did not exhibit this phenotype, and instead consistently doubled at the same rate over 10 passages in culture (Figure 2A and Figure 7B). Whole genome sequencing of the rapidly-growing Ts19 cell line at late passage showed that it had unexpectedly gained an extra copy of chromosome 2 in addition to the trisomy of chromosome 19 (Figure 7A). In contrast, the slower-growing Ts19 cell line was found to maintain its initial karyotype (Figure 7B). Similarly, in a set of independent cell lines, we noted that one line of Ts19+LTa+PIK3CA^H1047R^ MEFs proliferated more rapidly than its euploid control, while Ts19+LTa+Vec and Ts19+LTa+Ras^V12^ cells proliferated at the same rate as similarly-transduced euploid MEFs (Figure S8). Karyotype analysis demonstrated that the Ts19+LTa+PIK3CA^H1047R^ cell line had also gained an additional copy of chromosome 2, while the other Ts19 cell lines had not (Figure S11A). In a replicate experiment on a further set of independently-derived cell lines, Ts19+LTa cells transduced with an empty vector or three different oncogenes, including PIK3CA^H1047R^, were found to proliferate at the same rate as equivalently-transduced euploid cell lines (Figure S8). Read-depth analysis at late passage revealed a variety of karyotypes in these cell lines (Figure S11B). While three cell lines had gained extra copies of mChr2, this gain was accompanied by other alterations, including the gains of mChr1, mChr6, mChr15, mChr18, and the loss of mChr14. Thus, in every experiment in which Ts19 cells were found to proliferate more rapidly than euploid cells, their karyotype was found to be +mChr2+mChr19, while in seven other experiments this karyotype was not observed. These results suggest that the gain of mChr2 and mChr19, in the absence of other karyotypic changes, may enhance the proliferative capacity of cells expressing LTa.

**Figure 7.**
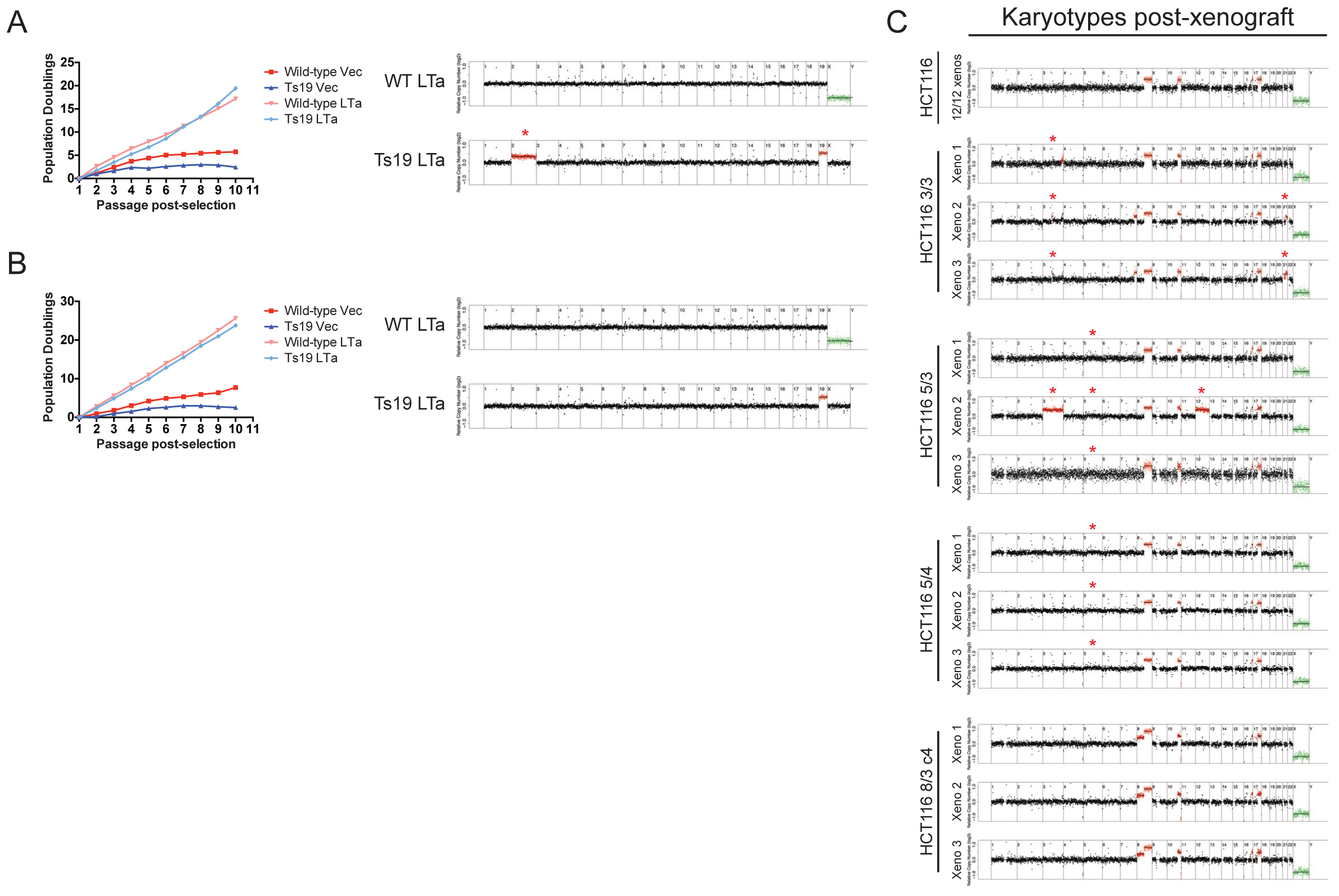
Karyotype evolution correlates with enhanced growth in aneuploid cell lines. (A) One line of Ts19 MEFs and a matched euploid control cell line were transduced with LTa or with an empty vector. Sequence analysis at passage 7 revealed an extra copy of chromosome 2 (indicated with an asterisk) in the Ts19 line. Note that this growth curve is also displayed in Figure S4, and is reproduced here for reference. (B) An independently-derived pair of WT and Ts19 cell lines were transduced as above. Sequence analysis at passage 7 revealed that the initial karyotypes of these cell lines had been maintained. Note that this growth curve is also displayed in Figure 2A, and is reproduced here for reference. (C) Xenografts from figure 6 were extracted, digested with trypsin, and then plated on plastic. Low-pass whole genome sequencing revealed that 12 HCT116 xenografts and 3 HCT116 8/3 c4 xenografts maintained their initial karyotypes. However, all HCT116 3/3, 5/3, and 5/4 xenografts lost their initial trisomies or tetrasomies during *in vivo* growth, and several lines displayed additional chromosomal copy number alterations. Deviations from each cell line’s initial karyotype are indicated with an asterisk.

To expand our analysis beyond Ts19, we next analyzed a Ts1+p53dd+Ras^V12^ cell line. These cells initially doubled every ~50 hours. After 10 passages in culture, the trisomic cell line was observed to double every ~27 hours, a rate indistinguishable from p53dd+Ras^V12^-transduced wild-type cells (Figure 3A). Whole genome sequencing at early passage demonstrated that the euploid and trisomic cell lines maintained their initial karyotypes following transduction. However, at passage 10, a second round of genome sequencing demonstrated that the rapidly-growing Ts1+p53dd+Ras^V12^ cell line had lost the extra copy of chromosome 1, and instead displayed several other chromosome gains and losses, which were consistent with trisomies and pentasomies in a tetraploid population (Figure S12). The Ts1+p53dd+Vec line, which continued to proliferate at a very low rate, maintained an extra copy of chromosome 1 at late passage. Thus, improved growth of an aneuploid cell line may also result from chromosome loss and/or tetraploidization.

To determine whether similar adaptations occur *in vivo*, we re-derived cell lines from 12 HCT116 xenografts and from 3 HCT116 3/3, 5/3, 5/4, and 8/3 c4 xenografts. Following two passages in culture to deplete stromal cell contamination, each re-derived cell line was subjected to whole-genome sequencing. The euploid cell lines were chromosomally stable, and 12 out of 12 re-derived lines had karyotypes that were indistinguishable from the pre-xenograft cell line (Figure 7C). However, every HCT116 3/3, 5/3 and 5/4 cell line was found to have lost its original trisomic or tetrasomic chromosome. 3 out of these 9 aneuploid lines exhibited other chromosomal alterations: one cell line initially trisomic for chromosome 5 gained an extra copy of chromosomes 3 and 12, while two cell lines initially trisomic for chromosome 3 gained an extra copy of chromosome 21. HCT116 8/3 c4, which proliferated at nearly the same rate as the parental line, maintained its trisomy when grown as a xenograft, and did not exhibit any further chromosomal alterations. These results indicate that growth-inhibitory aneuploidies are selected against during *in vivo* tumor formation, while aneuploidies that are neutral can escape negative selection.

## Discussion

Aneuploidy is a nearly-universal feature of human cancers. However, despite its frequent occurrence, we have found that in carefully-controlled experiments, aneuploid cells exhibit reduced tumorigenicity relative to genetically-matched euploid cells. Gain of 6 different chromosomes tested so far (mChr1, mChr13, mChr16, mChr19, hChr3, and hChr5) impedes several measures of tumorigenic capacity, while gain of only one chromosome (hChr8) has nearly-neutral consequences. Moreover, the activation of several oncogenic pathways (HRAS, BRAF, PIK3CA, and MYC) as well as the ablation of p53 and RB function are insufficient to overcome the fitness penalty induced by aneuploidy. While certain oncogene cocktails, particularly those that include LTa or Ras^V12^, result in a partial and variable suppression of the aneuploidy-induced proliferation delay, trisomic cells transduced with multiple oncogenes exhibit consistent and severe defects in focus formation, anchorage-independent growth, and growth as xenografts. We failed to detect any conditions in which aneuploidy promotes the transformation of primary cells, synergized with an oncogenic mutation, or otherwise contributed to tumorigenesis. Instead, we have found that many aneuploidies are actively selected against during growth *in vitro* or *in vivo*.

While these results are unexpected, they are consistent with much of what has been learned about the effects of whole-chromosome aneuploidy on normal cell physiology. Aneuploid chromosomes are transcribed and translated proportional to their copy number (Dephoure et al., 2014; Stingele et al., 2012; Torres et al., 2007, 2010), which can lead to stoichiometric imbalances in endogenous proteins and protein complexes (Oromendia et al., 2012; Sheltzer and Amon, 2011). To compensate, cells rely on a set of protein quality control mechanisms, including the HSF1/HSP90 folding pathway (Donnelly et al., 2014; Oromendia et al., 2012), autophagy, and proteasomal degradation (Dephoure et al., 2014; Torres et al., 2010). The energetic cost of expressing, folding, and turning over excess proteins, as well as the downstream consequences of unmitigated protein imbalances, impose a significant fitness cost on the cell. The severity of aneuploid phenotypes is generally proportional to the degree of aneuploidy, and indeed, we found that transformed cells trisomic for mouse chromosome 1 grew much more slowly than cells trisomic for mouse chromosome 19. Moreover, while studies of mouse models of Down syndrome have identified certain genes whose triplication may have tumor-protective effects (Baek et al., 2009; Reynolds et al., 2010; Sussan et al., 2008), our results suggest that whole-chromosome aneuploidy itself can function as a powerful tumor suppressor.

The molecular pathway(s) that cause slow cell division and tumor suppression in response to aneuploidy remains an area of active investigation. We found that aneuploidy commonly induces senescence in both primary and transformed cells, as has previously been reported in cells that display chromosomal instability (Baker et al., 2004; Lentini et al., 2012). How aneuploidy induces senescence is an important question that remains to be addressed. However, our findings that HCT116 5/3 and E1a-transduced trisomic MEFs exhibit both minimal senescence and poor proliferation suggest that other factors contribute to the growth defect of aneuploid cells as well.

If aneuploidy can function as a tumor suppressor, why is aneuploidy such a frequent occurrence in cancer? Several possibilities remain. First, our study has specifically examined the consequences of single-chromosome trisomies and tetrasomies. This degree of aneuploidy is frequently observed in early-stage lesions in a variety of tumor types (Balaban et al., 1986; Di Capua Sacoto et al., 2011; Lai et al., 2007; Magnani et al., 1994; El-Rifai et al., 2000). It could be the case that these low levels of aneuploidy are in fact tumor-protective, while complex karyotypes or the multiple chromosome gains found in more advanced malignancies are tumor-promoting. Monosomies could also have oncogenic consequences that trisomies and tetrasomies lack. Novel genetic tools will be required to generate and study these types of defined karyotype changes without inducing gross CIN. Nonetheless, our results argue that the simple aneuploidies found in pre-malignant cells may not be oncogenic.

Alternately, aneuploidy could function predominantly as a tumor suppressor, yet still exhibit tumor-promoting effects under very rare circumstances that involve specific cell types, chromosomes, or mutational backgrounds. For example, trisomy of chromosome 21 predisposes individuals to leukemia (Seewald et al., 2012), and gain of chromosome 21 is a common occurrence in sporadic leukemia (Loncarevic et al., 1999; Ozery-Flato et al., 2011), but trisomy of chromosome 21 appears to protect against the development of many other cancer types, including breast, lung, and prostate cancers (Nižetić and Groet, 2012). Thus, while the nine aneuploid lines that we examined hinder or are neutral with regard to tumor growth, it is conceivable that a wider survey of aneuploidies, oncogenes, or cell types would reveal unusual cases in which aneuploidy provides a growth advantage. Under natural selection in a tumor, rare growth-promoting aneuploidies could rise to clonal levels while a large number of growth-inhibitory aneuploidies are continually selected against. Finally, chromosome missegregation, rather than aneuploidy *per se*, could be a crucial driver of tumorigenesis. Lagging chromosomes can be damaged during anaphase (Janssen et al., 2011) or dramatically altered following encapsulation in a micronucleus (Crasta et al., 2012); missegregation-induced DNA damage could therefore promote transformation while any subsequent aneuploidy exists mainly as a “passenger” mutation. Similarly, reductive mitoses from a tetraploid intermediate may produce tumor-initiating gross chromosomal rearrangements with a large number of aneuploid chromosome “passengers” (Fujiwara et al., 2005). Thus, while we have found that single-chromosome trisomies function as tumor suppressors, other types of aneuploidy or other routes of generating aneuploid cells could have oncogenic consequences.

Tumor cells may also adapt to suppress certain adverse effects of aneuploidy, and our results demonstrate one potential mechanism by which this can occur. Following serial passaging or growth *in vivo*, aneuploid cells frequently exhibit further karyotypic alterations. Remarkably, these changes correlate with improved growth. Aneuploid cells can revert to euploidy by losing their extra chromosomes, and we have found that this is a common occurrence when the extra chromosome(s) induces a significant growth disadvantage. Alternately, cells can acquire other chromosome copy number changes, including both chromosome gains and losses. We propose that aneuploidy + oncogene “sweet spots” exist in which the detrimental effects of aneuploidy are neutralized while a pro-proliferation phenotype is uncovered. In particular, the gain of mChr2 was found to correlate with enhanced growth in multiple independent experiments with Ts19. It may be the case that the gain of mChr2 and mChr19 in cells expressing LTa is one such “sweet spot,” and that these aneuploidy sweet spots represent frequently-observed karyotypes in cancer. Consistent with this notion, many distinct chromosome copy number alterations are observed together in the same tumors more often than expected by chance (Ozery-Flato et al., 2011). For instance, tumors that have gained an extra copy of hChr7 are significantly more likely to have also gained an extra copy of hChr17, while loss of hChr7 is correlated with loss of hChr17 (Ozery-Flato et al., 2011). Such changes could potentially function to maintain stoichiometry in key protein complexes while also promoting tumorigenic growth through as-yet undiscovered pathways. The karyotypic plasticity of aneuploid genomes that we have uncovered could drive the development of cancers that harbor these favorable karyotypic combinations.

## Materials and Methods

### MEF derivation, culture, and transduction

A Robertsonian breeding scheme was utilized to generate sibling-matched euploid and trisomic MEFs as described in Williams et al., 2008. Note that due to the extreme fitness defect caused by the gain of chromosome 1, to date we have only been able to generate a MEF line from a single Ts1 embryo. Therefore, only single replicates of experiments involving this trisomy are displayed.

MEFs were cultured in DMEM supplemented with 10% FBS, 2mM glutamine, and 100 U/ml penicillin and streptomycin. Cells were maintained at 37***°***C and 5% CO_2_ in a humidified environment. Cell counting was performed using the Cellometer Auto T4 system. Plasmids encoding oncogenes were obtained from Addgene (https://www.addgene.org/) and then transfected into the Phoenix-Eco cell line (Swift et al., 2001) using TransIT-LT1 (Mirus). Viral supernatants were collected 24, 48, and 72 hours post-transfection, and were applied to freshly-split passage 2 MEFs. Transduced cells were selected by FACS, or by the addition of puromycin (1.6 µg/ml), hygromycin (200 µg/ml), or G418 (1 mg/ml).

### Human colon cancer cell culture

Aneuploid cell lines derived from HCT116 cells were previously described in Stingele et al., 2012 and Donnelly et al., 2014. Cells were cultured in DMEM supplemented with 10% FBS, 2mM glutamine, and 100 U/ml penicillin and streptomycin. Cells were maintained at 37***°***C and 5% CO_2_ in a humidified environment.

### Low-pass whole genome sequencing

Sequencing reactions were performed at the MIT BioMicro Center. 50 ng of purified DNA from each cell line were prepared and barcoded using Nextera reagents (Illumina), and tagmented material was PCR amplified for seven cycles. Libraries were quantified using an AATI Fragment Analyzer before pooling. Libraries were sequenced (40bp read length) on an Illumina HiSeq2000. Reads were demultiplexed using custom scripts allowing single mismatches within the reference barcode.

Sequence reads were trimmed to 40 nucleotides and aligned to the mouse (mm9) or human (hg19) genomes using BWA (0.6.1) with default options (Li and Durbin, 2009). HMMcopy (0.1.1) was used to detect copy number alterations by estimating copy number in 500-kb bins controlling for mappability [downloaded from UCSC Genome Bioinformatics (http://hgdownload.cse.ucsc.edu/goldenPath/hg19/encodeDCC/wgEncodeMapability/ or http://hgdownload.cse.ucsc.edu/goldenPath/mm9/encodeDCC/wgEncodeMapability/)] and GC content (calculated by HMMcopy gcCounter)(Ha et al., 2012).

### Cell proliferation and tumorigenicity assays

For proliferation assays, MEFs and HCT116 cells were passaged using a modified 3T3 protocol (Todaro and Green, 1963). 3×10^5^ cells were plated in three wells of a 6-well plate, and cells were combined, counted, and re-plated at the same density every third day. For focus formation assays, 1000 cells (MEFs) or 200 cells (HCT116) were plated in triplicate on 10cm plates, and then allowed to grow for 10 (MEFs) or 14 days (HCT116). Subsequently, colonies were fixed with ice-cold 100% methanol for 10 minutes, and then stained with a solution of 0.5% crystal violet in 25% methanol for 10 minutes. For soft agar assays, a 1% base layer of Difco Agar Noble was prepared and then mixed with an equal amount of 2X DMEM. The solution (0.5% agar in 1X DMEM) was then added to each well of a 6-well plate and allowed to solidify. Subsequently, a top layer of 0.7% agar was prepared and mixed with an equal volume of a 2X solution of DMEM containing 10,000 cells (MEFs) or 2000 cells (HCT116) and added to the base layer in triplicate. The plates were incubated for 20 days at 37***°***C prior to imaging.

For xenograft studies, 5 ‐ 10 female, 5-week old Nu/J mice (Jackson Laboratory Stock 002019) were utilized for each experiment. Cells to be injected were harvested and concentrated to 10^7^ (MEFs) or 4x10^7^ (HCT116) cells/ml in PBS. 100µl of the solution was injected subcutaneously into the rear flanks of each mouse using a 25 gauge needle. Euploid and aneuploid cell lines were typically injected contralaterally, with the exception of experiments involving cell lines transduced with LTa and Ras^V12^, in which only one cell line was injected into each animal. Tumor dimensions were measured every third day using calipers, and tumor volumes were calculated using the formula 0.5 × A × B^2^, where A is the longer diameter and B is the shorter diameter. H&E staining of paraformaldehyde-fixed sections was performed according to standard methods. All animal studies and procedures were approved by the MIT Institutional Animal Care and Use Committee.

### β-galactosidase staining

5000 cells of each cell line were plated in triplicate in a 48-well plate, allowed to attach overnight, and then stained using a Senescence Histochemical Staining Kit (Sigma-Aldritch). Cells were incubated in the X-gal solution overnight at 37***°***C prior to imaging on a Nikon Eclipse TE2000.

## Acknowledgments

We thank the Koch Institute Flow Cytometry facility for assistance with cell sorting. We thank the MIT BioMicro Center for performing sequencing reactions, and Charlie Whittaker and Jie Wu in the Koch Institute Bioinformatics Core for assistance with data analysis (NIH grant P30-CA14051). This work was supported by the National Institute of Health GM056800 to A.A and the Kathy and the Curt Marble Cancer Research Fund. J.S. was supported by a Whitaker Health Sciences Fund Fellowship and an MIT School of Science Fellowship in Cancer Research. A.A. is an investigator of the Howard Hughes Medical Institute and the Glenn Foundation for Biomedical Research.

## Supplemental Figure Legends

**Figure S1.**
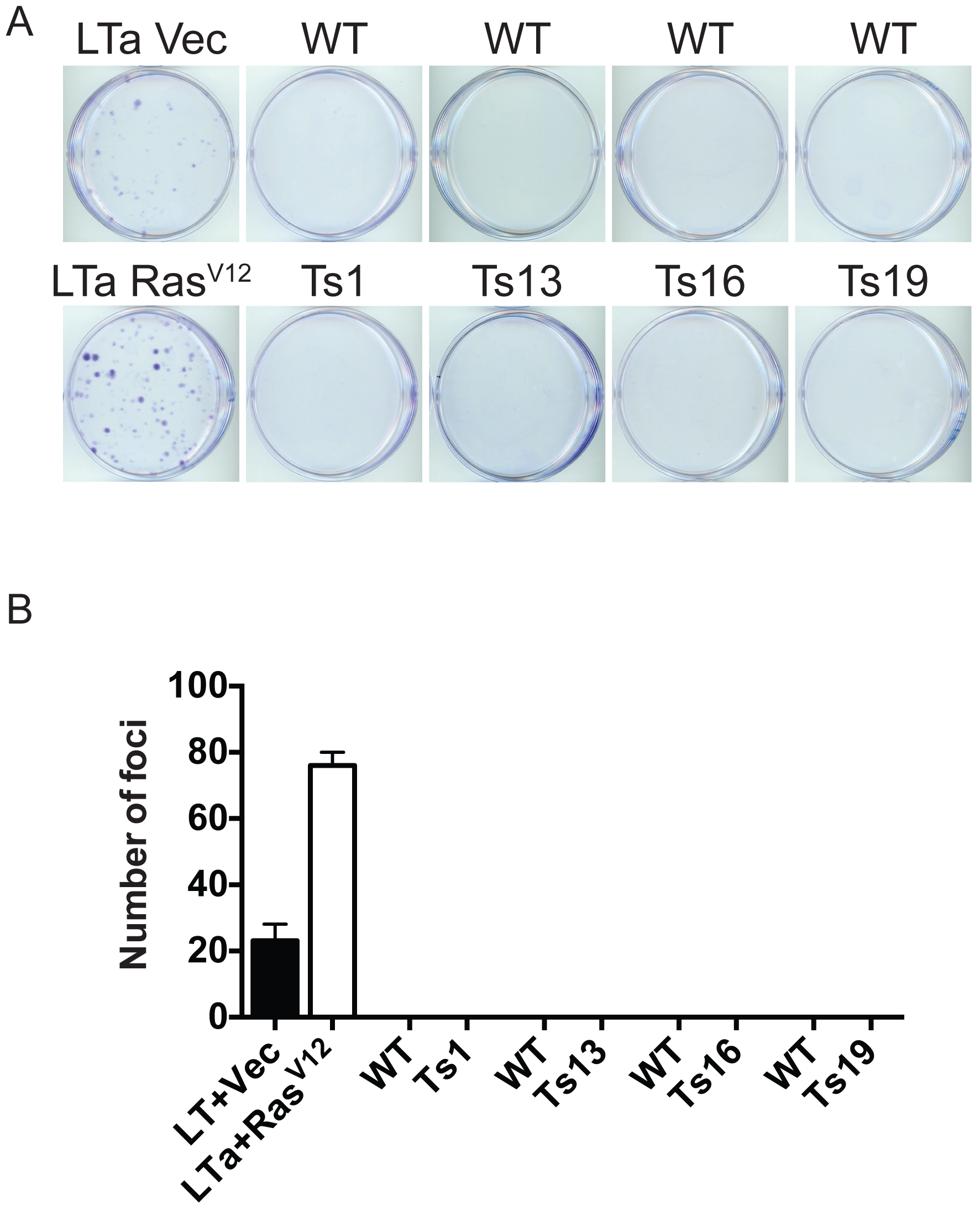
Primary aneuploid cells are non-clonogenic. (A) 1000 cells of the indicated cell lines were plated and grown for 10 days prior to staining with crystal violet. LTa-transduced MEFs are capable of forming colonies from single cells, but primary euploid and trisomic MEFs are non-clonogenic. (B) Quantification of (A).

**Figure S2.**
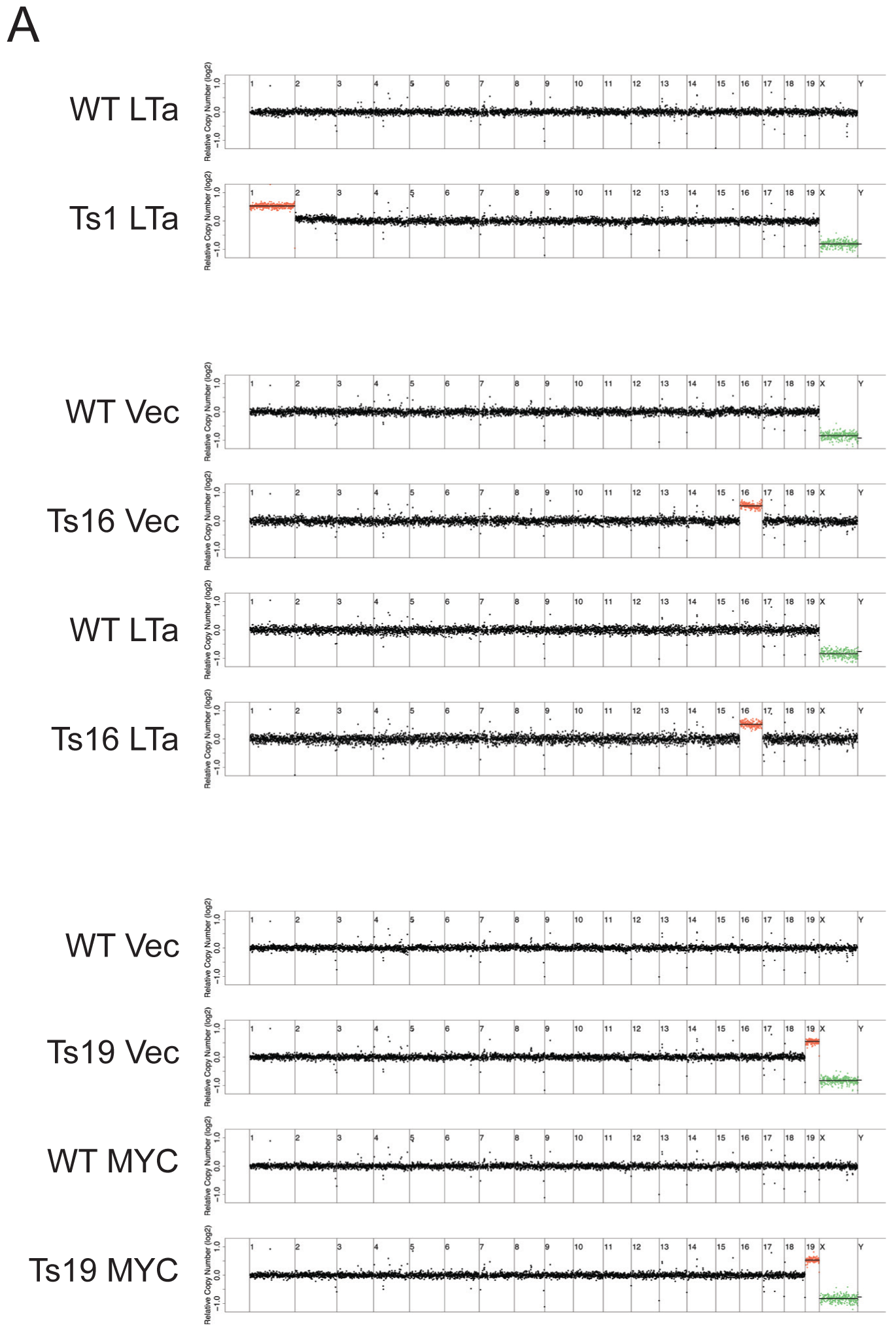
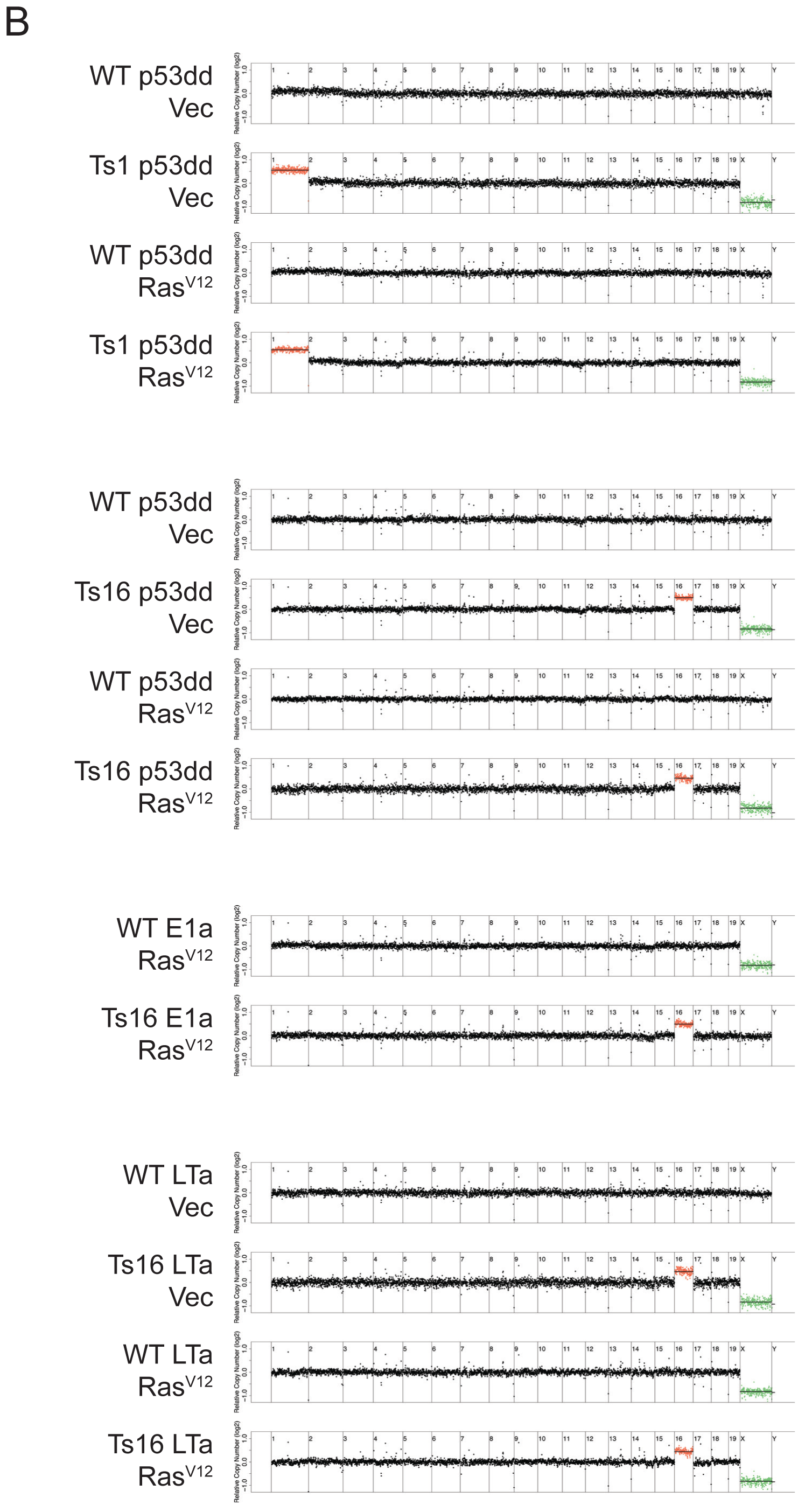
Trisomic karyotypes are maintained during retroviral transduction and selection. Karyotypes of MEF lines that were transduced with (A) one oncogene or (B) two oncogenes were determined by low-pass whole genome sequencing. Note that the sequence results from the Ts1+p53dd+Ras^V12^ and control cell lines are reproduced in Figure S12.

**Figure S3.**
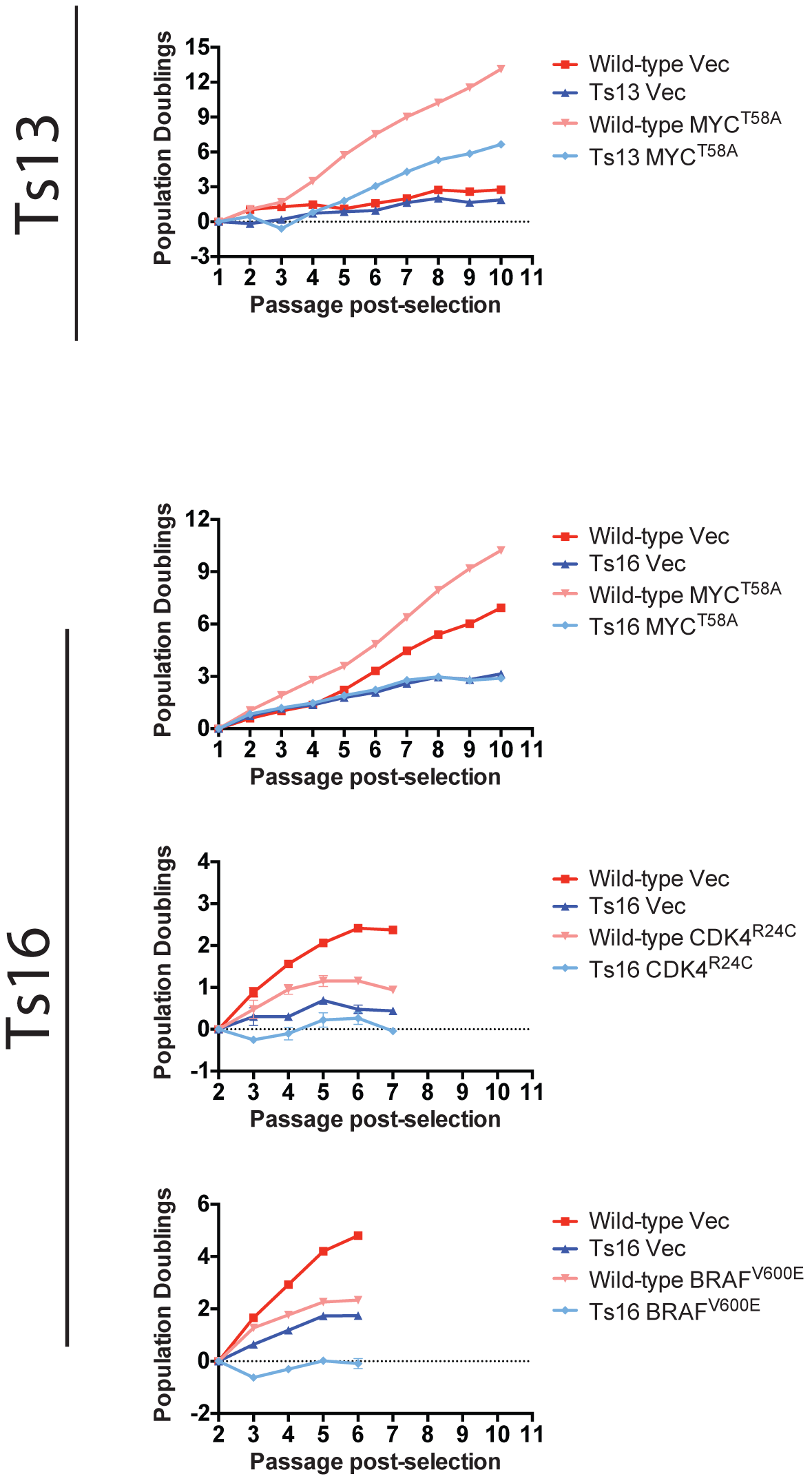
Effects of MYC^T58A^, BRAF^V600E^, and CDK^R24C^ on the proliferation of euploid and trisomic cells. Euploid and trisomic cell lines were transduced with plasmids harboring the indicated oncogene or a matched empty vector. Following selection, the cell lines were passaged every third day for up to 10 passages, and the cumulative population doublings were determined. Experiments with BRAF^V600E^, and CDK4^R24C^ were terminated prematurely as both euploid and trisomic cells senesced following transduction.

**Figure S4.**
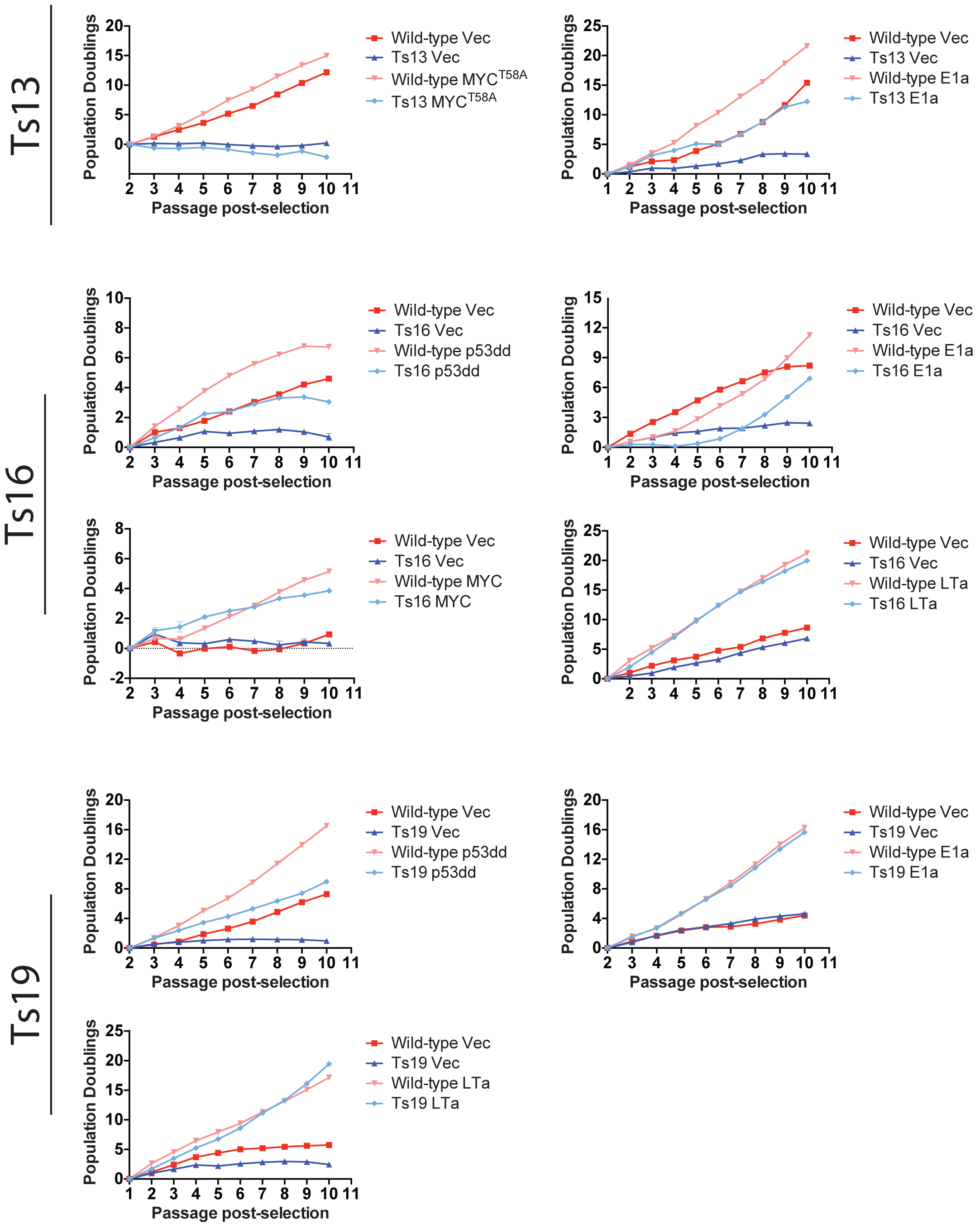
Replicate oncogene-transduction experiments on independently-derived cell lines. Nine single-oncogene transduction experiments were repeated in different, independently-derived cell lines. These lines were stably transduced with plasmids harboring the indicated oncogene(s) or a matched empty vector, and then passaged every third day for up to 10 passages. Some variability exists between replicates (e.g., compare Ts16+MYC in Figure S4 and Figure 2A), but no trisomy+oncogene combination was found to consistently outgrow a matched euploid line across multiple cell lines. Note that the panel displaying Ts19+LTa is reproduced in Figure 7.

**Figure S5.**
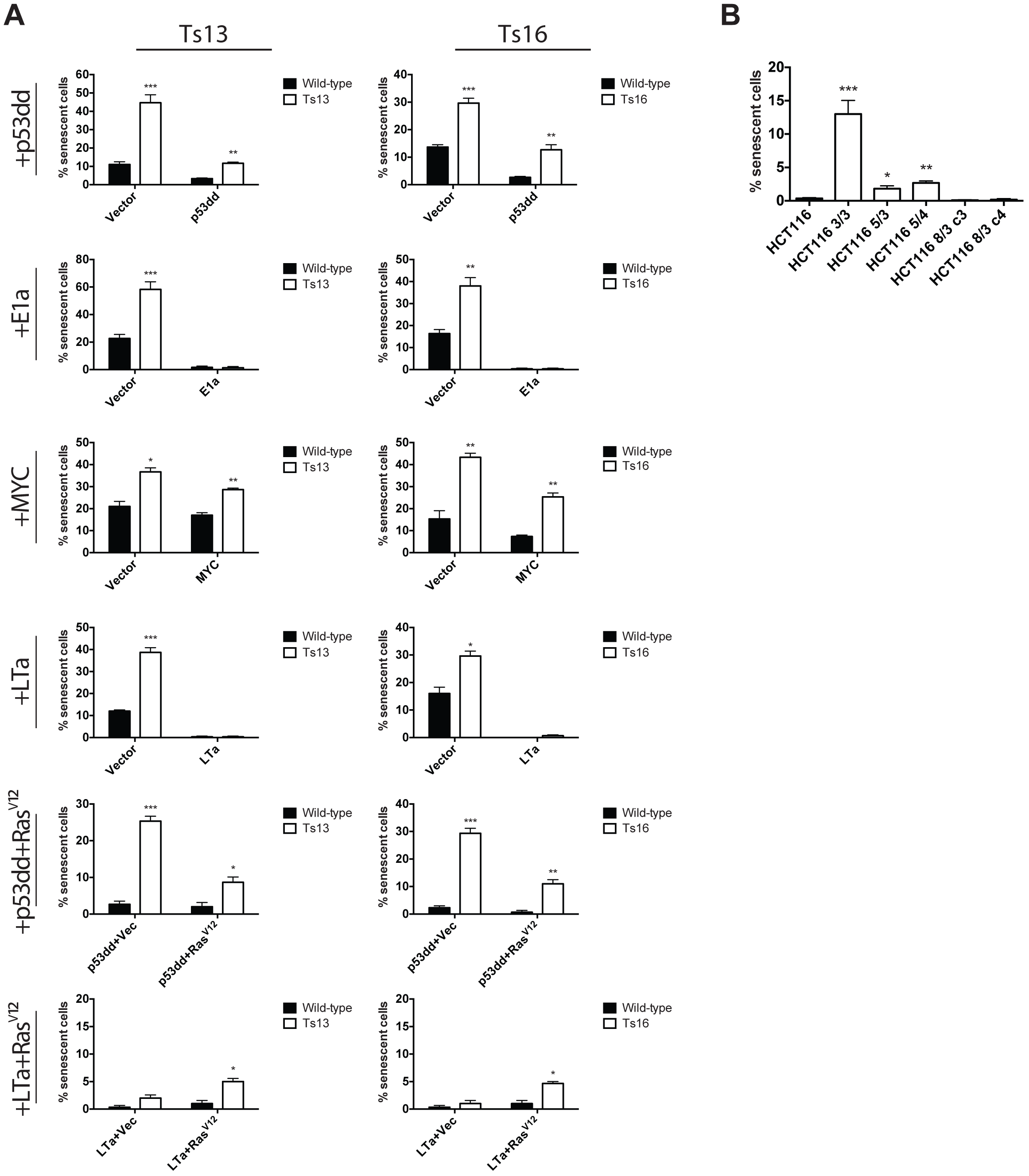
Aneuploid cells display elevated levels of senescence-associated β-galactosidase. At passage 6-8 in culture, the indicated cells were plated and stained for the expression of β-galactosidase. At least 200 cells in three wells were counted for each experiment. *, p<.05; **, p<.005; ***, p<.0005 (Student’s t test).

**Figure S6.**
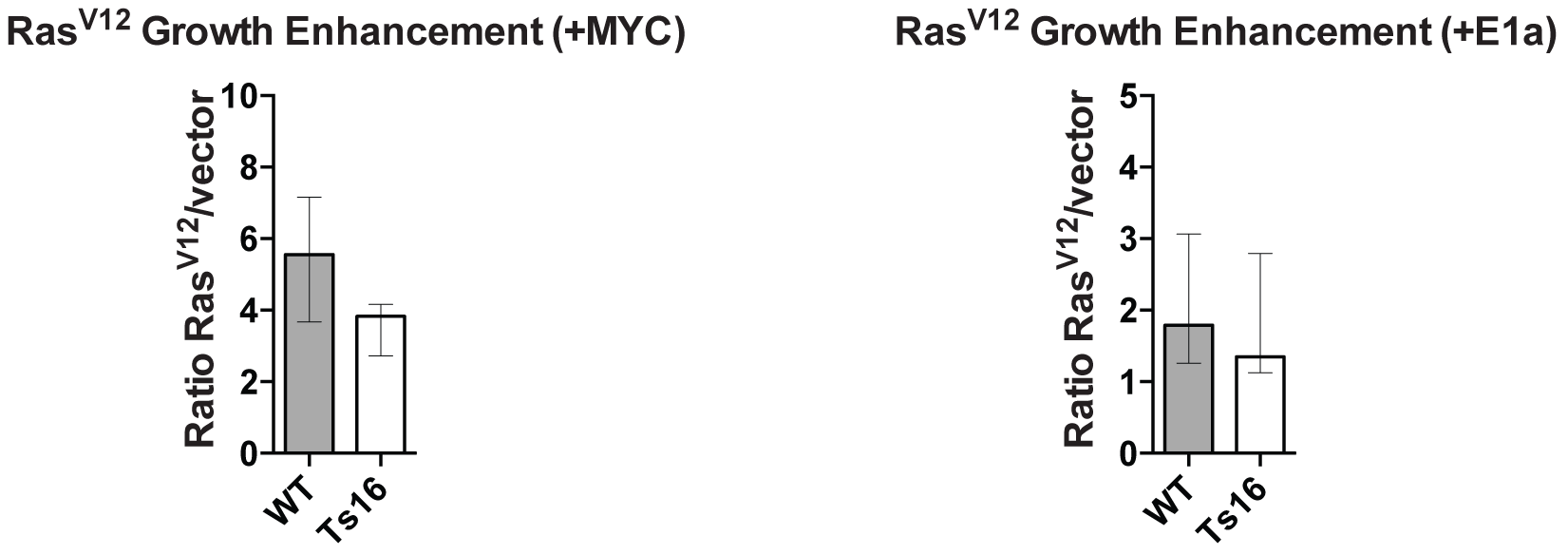
Relative growth enhancement conferred by oncogenes in euploid and aneuploid cell lines. The number of cells recovered from Ras^V12^-transduced MEFs was divided by the number of cells recovered from vector-transduced MEFs at every passage. Bar graphs display the median ratios and the interquartile ranges. * p<.05; *** p<.0005 (Wilcoxon rank-sum test).

**Figure S7.**
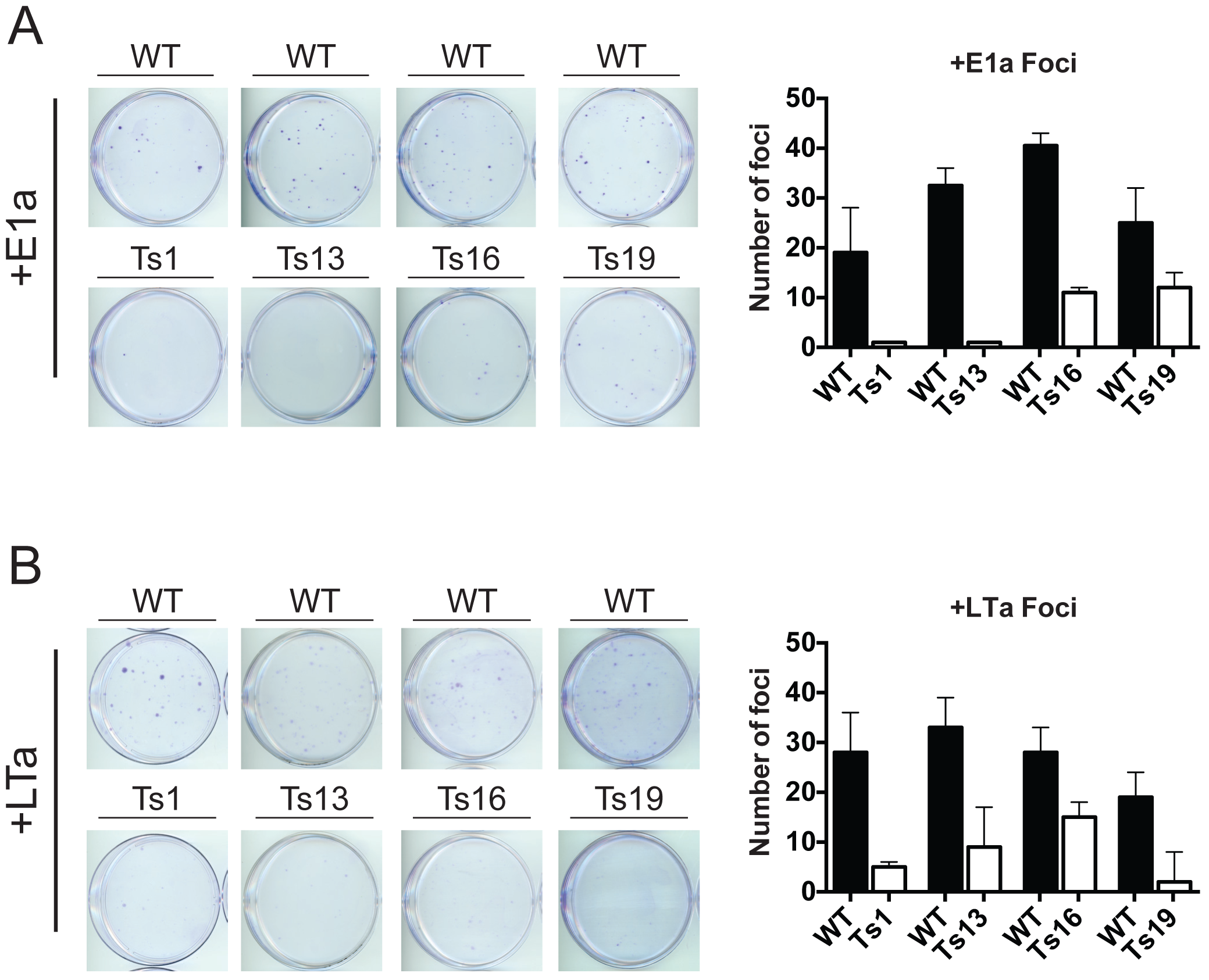
Oncogene-transduced aneuploid cell lines exhibit reduced clonogenicity. (A and B) 1000 cells of the indicated cell lines were plated and then allowed to grow for 10 days before being stained with crystal violet. Representative plates are shown on the left, while average colony counts are displayed on the right. In each experiment, the trisomic cell lines were found to exhibit a significantly reduced focus formation ability relative to the matched euploid cell line (p<.01, Student’s t test).

**Figure S8.**
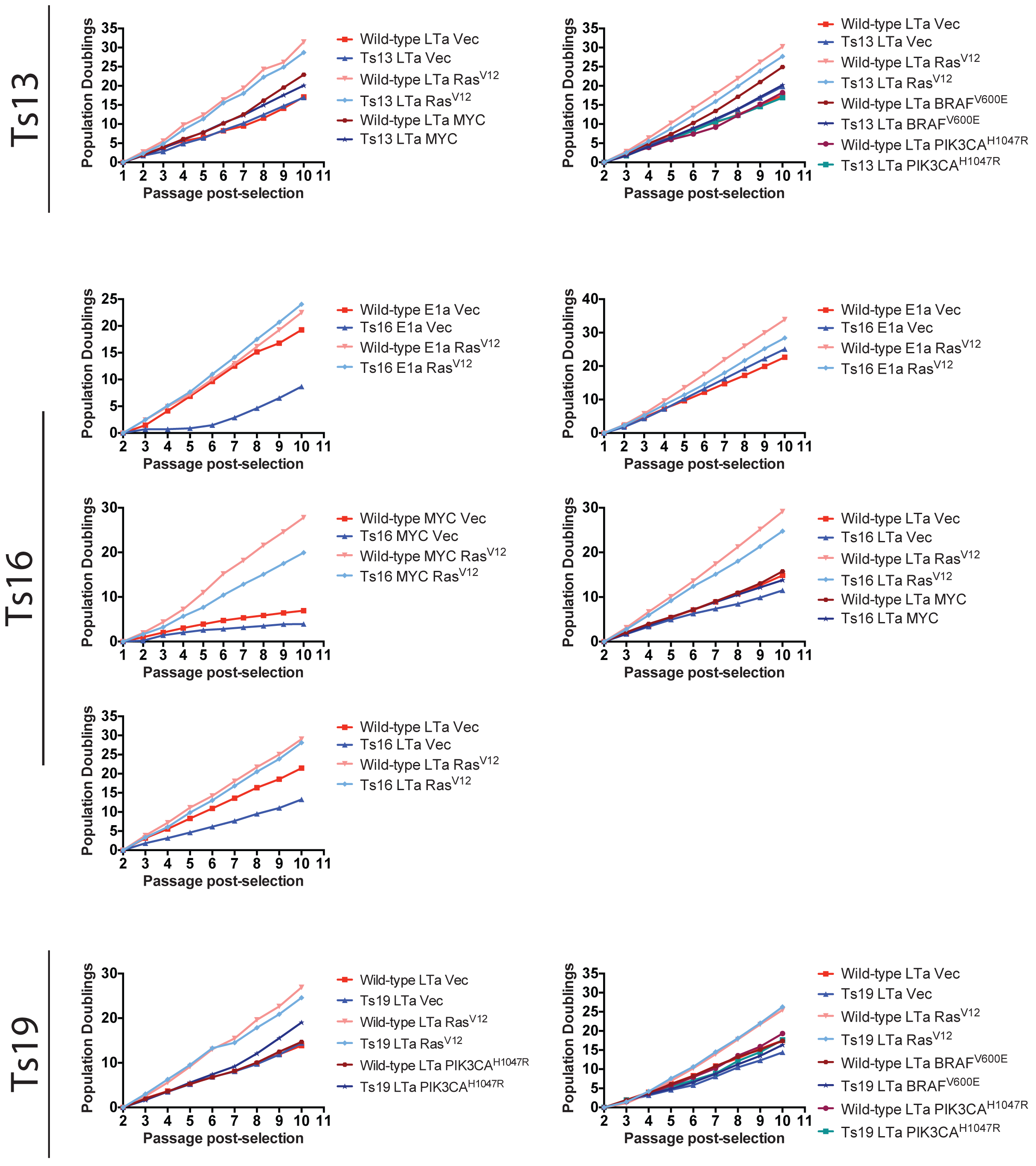
Additional oncogene cocktails and replicate experiments in trisomic MEFs. Euploid and trisomic cell lines were stably transduced with plasmids harboring the indicated oncogene or a matched empty vector. Following selection, the cell lines were passaged every third day for up to 10 passages, and the cumulative population doublings over the course of each experiment are displayed. Note that one replicate of Ts19+LTa+Vec and +Ras^V12^ (right panel) is also presented in Figure 3A; these growth assays were conducted in parallel with growth assays in cells that harbor BRAF^V600E^ or PIK3CA^H1047R^ and are therefore shown here for comparison. Both Ts19+LTa experiments are also reproduced in Figure S11.

**Figure S9.**
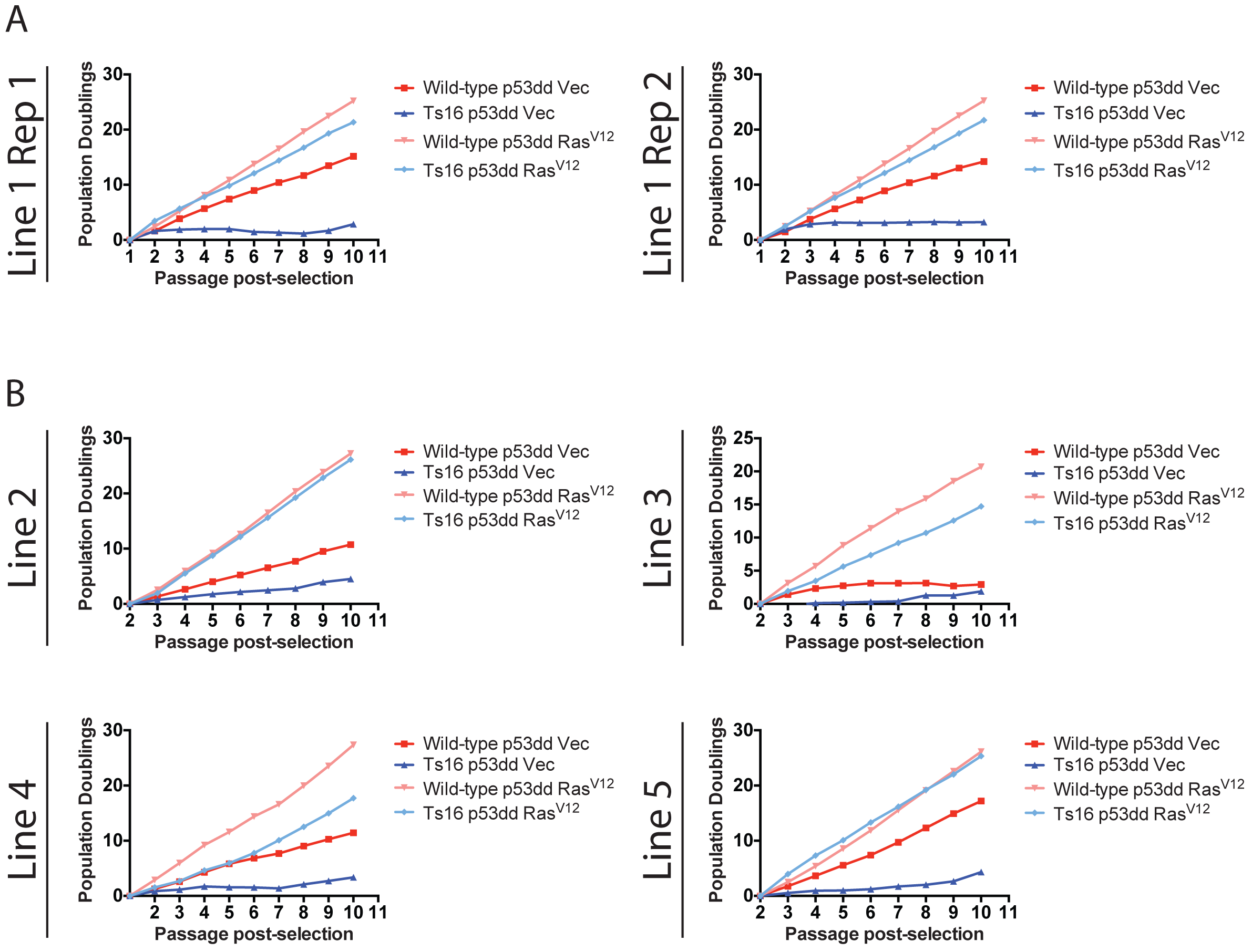
Replicate growth curves of the same or of independently-derived Ts16 cells that express p53dd and Ras^V12^. (A) The same WT and Ts16 MEF lines were stably transduced with plasmids harboring the indicated oncogene or a matched empty vector. Following selection, the cell lines were passaged every third day for up to 10 passages, and the cumulative population doublings over the course of each experiment are displayed. Note that Line 1 Rep 1 is also displayed in Figure 2A. (B) Four independently derived pairs of WT or Ts16 MEFs were transduced and passaged as described above.

**Figure S10.**
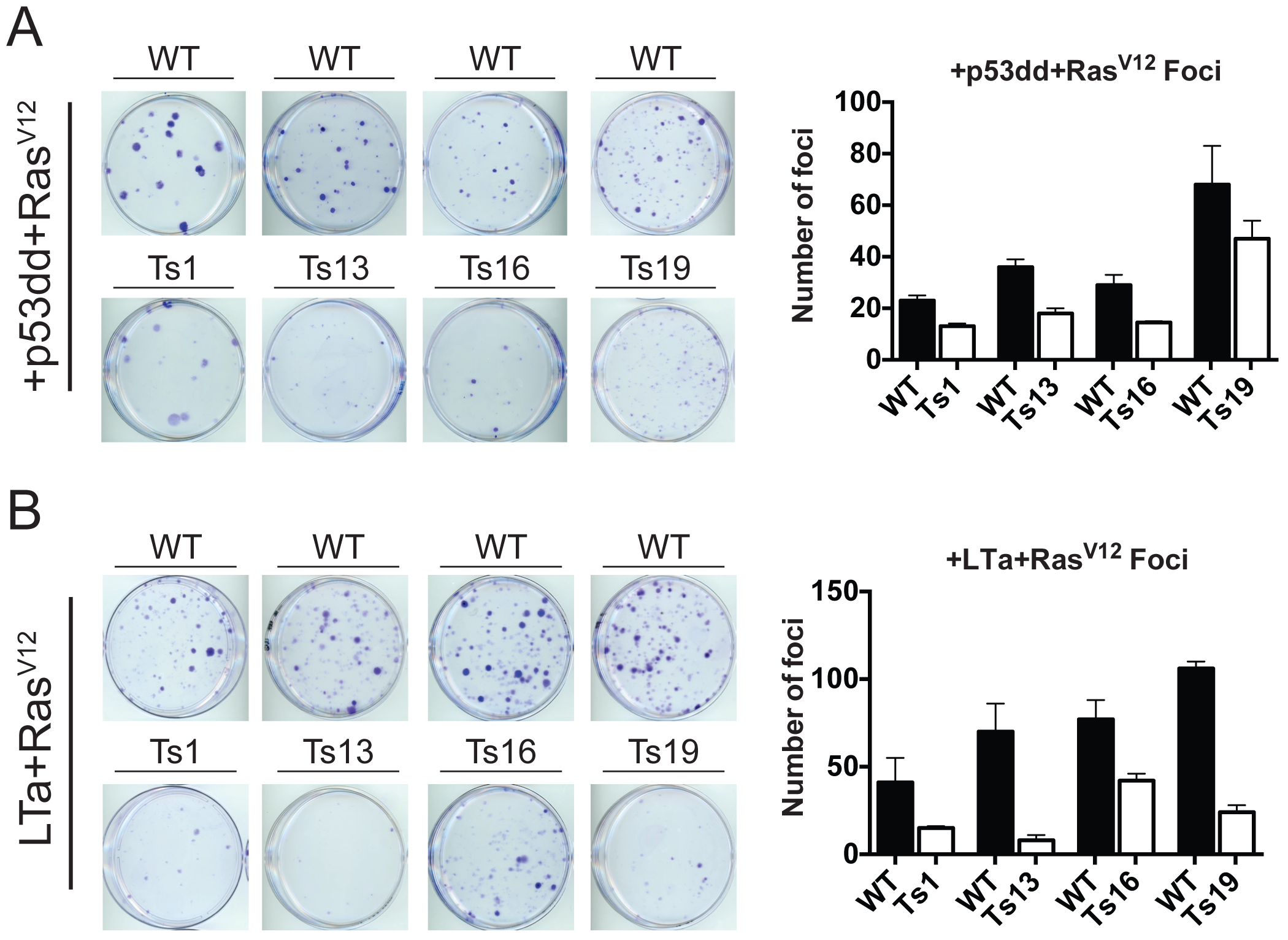
Transformed aneuploid cells display poor clonogenicity and a reduced ability to form colonies in soft agar. (A and B) 1000 cells of the indicated cell lines were plated and then allowed to grow for 10 days before being stained with crystal violet. For each comparison, the euploid MEFs formed more foci than the trisomic MEFs (p<.01, Student’s t test).

**Figure S11.**
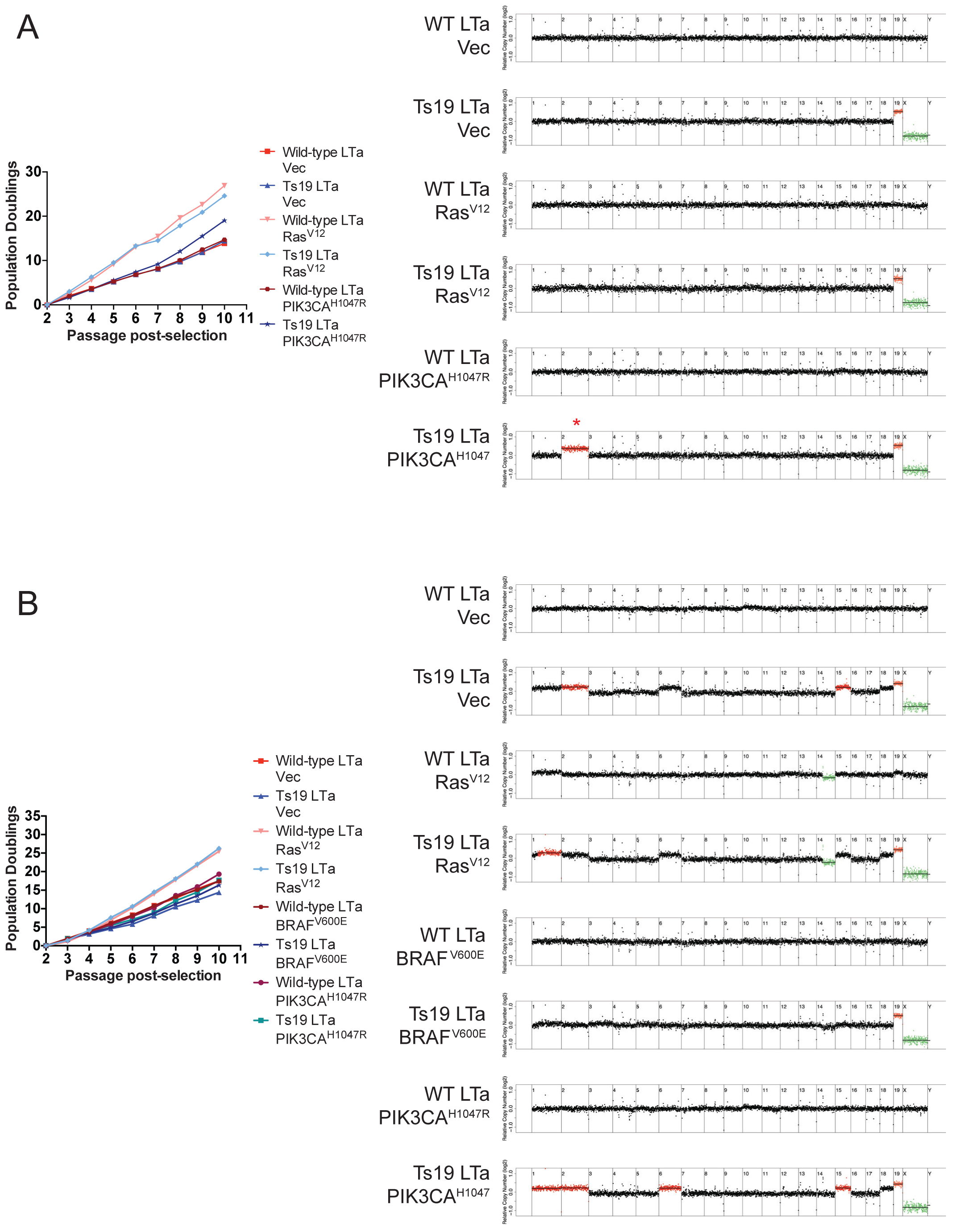
A rapidly-growing Ts19 line has gained mChr2 in the absence of other karyotype alterations. (A) A Ts19 cell line and a matched euploid control cell line were transduced with Large T, and then transduced a second time with an empty vector, Ras^V12^, or PIK3CA^H1047R^. The Ts19+LTa+PIK3CA^H1047R^ cell line grew more rapidly an equivalently-transduced euploid line. Sequencing at passage 10 revealed that the Ts19+LTa+PIK3CA^H1047R^ line had acquired an extra copy of chromosome 2, while the other cell lines showed no other deviations from the expected karyotypes. (B) An independently-derived Ts19 cell line and a matched euploid control cell line were transduced with LTa, and then transduced a second time with an empty vector, Ras^V12^, BRAF^V600E^, or PIK3CA^H1047R^. All euploid/Ts19 pairs grew at approximately the same rates over the course of the experiment. Read depth analysis at passage 10 revealed several karyotypic alterations, including a gain of mChr2. Note that the growth curves displayed in A and B are also presented in Figure 3A and Figure S8.

**Figure S12.**
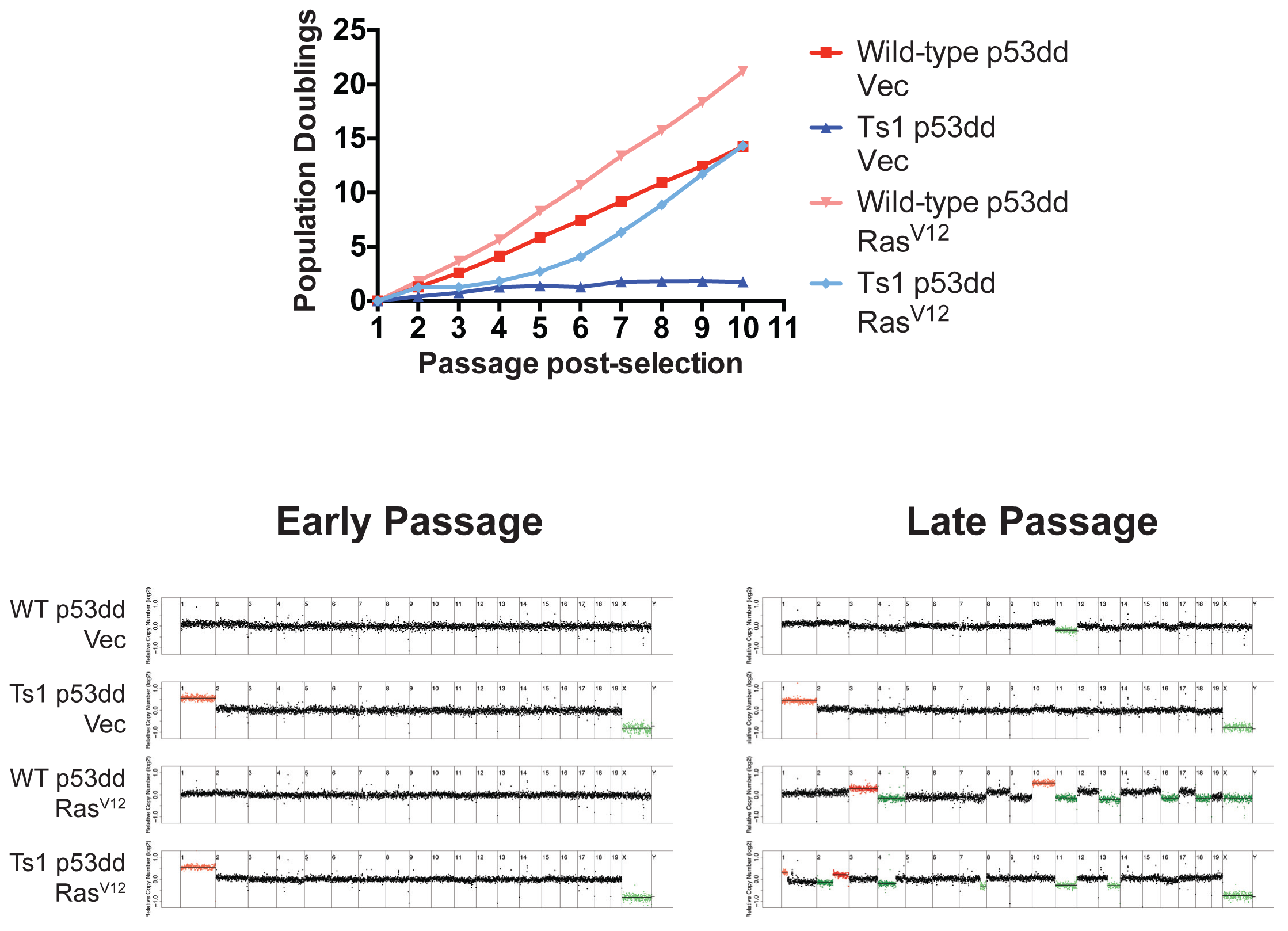
A rapidly-growing Ts1 line exhibits several karyotype changes. A Ts1 cell line and a matched euploid control line were transduced with p53dd and then transduced a second time with an empty vector or with Ras^V12^. The Ts1 MEFs initially grew poorly, then evolved to grow at a rate indistinguishable from that of wild-type cells. Whole-genome sequencing at passage 2 revealed that the slow growing line maintained its initial karyotype, while whole-genome sequencing at passage 10 revealed that the rapidly growing line had lost the trisomy of mChr1, and displayed several further chromosomal gains and losses. Note that this growth curve is also displayed in Figure 3A, and the early-passage karyotype analysis is also displayed in Figure S2B.

**Table S1.** Oncogenes present on mouse chromosomes 1,13,16, and 19.t

| **Chr.** | **Gene Symbol** | **Name** |
| --- | --- | --- |
| 1 | ELK4 | ELK4, ETS-domain protein (SRF accessory protein 1) |
| 1 | FCGR2B | Fc fragment of IgG, low affinity IIb, receptor for (CD32) |
| 1 | H3F3A | H3 histone, family 3A |
| 1 | MDM4 | Mdm4 p53 binding protein homolog |
| 1 | PBX1 | pre-B-cell leukemia transcription factor 1 |
| 1 | SLC45A3 | solute carrier family 45, member 3 |
| 1 | TPR | translocated promoter region |
| 1 | CREB1 | cAMP responsive element binding protein 1 |
| 1 | FEV | FEV protein - (HSRNAFEV) |
| 1 | IDH1 | isocitrate dehydrogenase 1 (NADP+), soluble |
| 1 | PAX3 | paired box gene 3 |
| 1 | SF3B1 | splicing factor 3b, subunit 1, 155kDa |
| 1 | NCOA2 | nuclear receptor coactivator 2 (TIF2) |
| 1 | TCEA1 | transcription elongation factor A (SII), 1 |
| 1 | BCL2 | B-cell CLL/lymphoma 2 |
| 1 | ABL2 | c-abl oncogene 2, non-receptor tyrosine kinase |
| 1 | PMX1 | paired mesoderm homeo box 1 |
| 1 | ACSL3 | acyl-CoA synthetase long-chain family member 3 |
| 1 | ATIC | 5-aminoimidazole-4-carboxamide ribonucleotide formyltransferase/IMP cyclohydrolase |
| 1 | CMKOR1 | chemokine orphan receptor 1 |
| 1 | LAF4 | lymphoid nuclear protein related to AF4 |
| 1 | FVT1 | follicular lymphoma variant translocation 1 |
| 1 | KDSR | 3-ketodihydrosphingosine reductase [Source:MGI Symbol;Acc:MGI:1918000] |
| 1 | LHX4 | LIM homeobox protein 4 [Source:MGI Symbol;Acc:MGI:101776] |
| 1 | FCGR3 | Fc receptor, IgG, low affinity III [Source:MGI Symbol;Acc:MGI:95500] |
| 1 | CSNK2A1 | casein kinase 2, alpha 1 polypeptide [Source:MGI Symbol;Acc:MGI:88543] |
| 13 | FGFR4 | fibroblast growth factor receptor 4 |
| 13 | IL6ST | interleukin 6 signal transducer (gp130, oncostatin M receptor) |
| 13 | NSD1 | nuclear receptor binding SET domain protein 1 |
| 13 | TERT | telomerase reverse transcriptase |
| 13 | DEK | DEK oncogene (DNA binding) |
| 13 | IRF4 | interferon regulatory factor 4 |
| 13 | TRIM27 | tripartite motif-containing 27 |
| 13 | OMD | osteomodulin |
| 13 | HIST1H3B | histone cluster 1, H3b |
| 13 | HIST1H4I | histone 1, H4i (H4FM) |
| 13 | SYK | spleen tyrosine kinase |
| 13 | NUTM2A | NUT family member 2A |
| 13 | NUTM2B | NUT family member 2B |
| 13 | NET1 | neuroepithelial cell transforming gene 1 [Source:MGI Symbol;Acc:MGI:1927138] |
| 13 | RASA1 | RAS p21 protein activator 1 [Source:MGI Symbol;Acc:MGI:97860] |
| **13** | SPZ1 | Spermatogenic leucine zipper protein 1 (BHLH-Zip transcription factor SPZ1) (Spermatogenic Zip 1) |
| **16** | BCL6 | B-cell CLL/lymphoma 6 |
| **16** | EIF4A2 | eukaryotic translation initiation factor 4A, isoform 2 |
| **16** | ETV5 | ets variant gene 5 |
| **16** | LPP | LIM domain containing preferred translocation partner in lipoma |
| **16** | TFG | TRK-fused gene |
| **16** | TFRC | transferrin receptor (p90, CD71) |
| **16** | CIITA | class II, major histocompatibility complex, transactivator |
| **16** | MYH11 | myosin, heavy polypeptide 11, smooth muscle |
| **16** | TNFRSF17 | tumor necrosis factor receptor superfamily, member 17 |
| **16** | ERG | v-ets erythroblastosis virus E26 oncogene like (avian) |
| **16** | OLIG2 | oligodendrocyte lineage transcription factor 2 (BHLHB1) |
| **16** | RUNX1 | runt-related transcription factor 1 (AML1) |
| **16** | TMPRSS2 | transmembrane protease, serine 2 |
| **16** | C16orf75 | chromosome 16 open reading frame 75 |
| **16** | RUNDC2A | RUN domain containing 2A |
| **16** | BCL5 | B-cell CLL/lymphoma 5 |
| **16** | PNUTL1 | peanut-like 1 (Drosophila) |
| **16** | IGL | immunoglobulin lambda locus |
| **16** | LITAF | LPS-induced TN factor [Source:MGI Symbol;Acc:MGI:1929512] |
| **16** | ETS2 | E26 avian leukemia oncogene 2, 3' domain [Source:MGI Symbol;Acc:MGI:95456] |
| **19** | CD274 | CD274 molecule |
| **19** | GNAQ | guanine nucleotide binding protein (G protein), q polypeptide |
| **19** | JAK2 | Janus kinase 2 |
| **19** | NFKB2 | nuclear factor of kappa light polypeptide gene enhancer in B-cells 2 (p49/p100) |
| **19** | NT5C2 | 5'-nucleotidase, cytosolic II |
| **19** | TCF7L2 | transcription factor 7-like 2 |
| **19** | TLX1 | T-cell leukemia, homeobox 1 (HOX11) |
| **19** | VTI1A | vesicle transport through interaction with t-SNAREs homolog 1A |
| **19** | MALAT1 | metastasis associated lung adenocarcinoma transcript 1 ( lnc-RNA; non-protein coding) |
| **19** | CD273 | programmed cell death 1 ligand 2 |
| **19** | KIAA1598 | KIAA1598 |
| **19** | MXI1 | Max interacting protein 1 [Source:MGI Symbol;Acc:MGI:97245] |

† Data are derived from the Sanger Cancer Genome Census and the Uniprot list of proto-oncogenes.

**Table S2.** Oncogenes present on human chromosomes 3, 5, and 8.t

| **Chr.** | **Gene Symbol** | **Name** |
| --- | --- | --- |
| 3 | BCL6 | B-cell CLL/lymphoma 6 |
| 3 | CTNNB1 | catenin (cadherin-associated protein), beta 1 |
| 3 | CCNL1 | Cyclin-L1 (Cyclin-L) |
| 3 | DCUN1D1 | DCN1-like protein 1 (DCUN1 domain-containing protein 1) (Defective in cullin neddylation protein 1-like protein 1) (Squamous cell carcinoma-related oncogene) |
| 3 | EVI1 | ecotropic viral integration site 1 |
| 3 | ETV5 | ets variant gene 5 |
| 3 | EIF4A2 | eukaryotic translation initiation factor 4A, isoform 2 |
| 3 | FOXL2 | forkhead box L2 |
| 3 | FOXP1 | forkhead box P1 |
| 3 | FHIT | fragile histidine triad gene |
| 3 | GATA2 | GATA binding protein 2 |
| 3 | GMPS | guanine monphosphate synthetase |
| 3 | LPP | LIM domain containing preferred translocation partner in lipoma |
| 3 | MECOM | MDS1 and EVI1 complex locus protein EVI1 (Ecotropic virus integration site 1 protein homolog) (EVI-1) |
| 3 | MITF | microphthalmia-associated transcription factor |
| 3 | MDS1 | myelodysplasia syndrome 1 |
| 3 | MYD88 | myeloid differentiation primary response gene (88) |
| 3 | MLF1 | myeloid leukemia factor 1 |
| 3 | NCKIPSD | NCK-interacting protein with SH3 domain (54 kDa VacA-interacting protein) (54 kDa vimentin-interacting protein) (VIP54) (90 kDa SH3 protein interacting with Nck) (AF3p21) (Dia-interacting protein 1) (DIP-1) (Diaphanous protein-interacting protein) (SH3 adapter protein SPIN90) (WASP-interacting SH3-domain protein) (WISH) (Wiskott-Aldrich syndrome protein-interacting protein) |
| 3 | PPARG | peroxisome proliferative activated receptor, gamma |
| 3 | PIK3CA | phosphoinositide-3-kinase, catalytic, alpha polypeptide |
| 3 | PRKCI | Protein kinase C iota type (EC 2.7.11.13) (Atypical protein kinase C-lambda/iota) (PRKC-lambda/iota) (aPKC-lambda/iota) (nPKC-iota) |
| 3 | RHOA | ras homolog family member A |
| 3 | RARB | Retinoic acid receptor beta (RAR-beta) (HBV-activated protein) (Nuclear receptor subfamily 1 group B member 2) (RAR-epsilon) |
| 3 | RPN1 | ribophorin I |
| 3 | AF3p21 | SH3 protein interacting with Nck, 90 kDa (ALL1 fused gene from 3p21) |
| 3 | SRGAP3 | SLIT-ROBO Rho GTPase activating protein 3 |
| 3 | SOX2 | SRY (sex determining region Y)-box 2 |
| 3 | TCTA | T-cell leukemia translocation-altered gene protein (T-cell leukemia translocation-associated gene protein) |
| 3 | TFRC | transferrin receptor (p90, CD71) |
| 3 | TFG | TRK-fused gene |
| 3 | USP4 | Ubiquitin carboxyl-terminal hydrolase 4 (EC 3.4.19.12) (Deubiquitinating enzyme 4) (Ubiquitin thioesterase 4) (Ubiquitin-specific-processing protease 4) |
|  |  | (Ubiquitous nuclear protein homolog) |
| 3 | RAF1 | v-raf-1 murine leukemia viral oncogene homolog 1 |
| 3 | WWTR1 | WW domain containing transcription regulator 1 |
| 3 | ZNF9 | zinc finger protein 9 (a cellular retroviral nucleic acid binding protein) |
| 5 | AFF4 | AF4/FMR2 family member 4 (ALL1-fused gene from chromosome 5q31 protein) (Protein AF-5q31) (Major CDK9 elongation factor-associated protein) |
| 5 | AF5q31 | ALL1 fused gene from 5q31 |
| 5 | CD74 | CD74 molecule, major histocompatibility complex, class II invariant chain |
| 5 | EBF1 | early B-cell factor 1 |
| 5 | FACL6 | fatty-acid-coenzyme A ligase, long-chain 6 |
| 5 | FGFR4 | fibroblast growth factor receptor 4 |
| 5 | GRAF | GTPase regulator associated with focal adhesion kinase pp125(FAK) |
| 5 | ITK | IL2-inducible T-cell kinase |
| 5 | IL6ST | interleukin 6 signal transducer (gp130, oncostatin M receptor) |
| 5 | IL7R | interleukin 7 receptor |
| 5 | LIFR | leukemia inhibitory factor receptor |
| 5 | CSF1R | Macrophage colony-stimulating factor 1 receptor (CSF-1 receptor) (CSF-1-R) (CSF-1R) (M-CSF-R) (EC 2.7.10.1) (Proto-oncogene c-Fms) (CD antigen CD115) |
| 5 | NSD1 | nuclear receptor binding SET domain protein 1 |
| 5 | NPM1 | nucleophosmin (nucleolar phosphoprotein B23, numatrin) |
| 5 | PDGFRB | platelet-derived growth factor receptor, beta polypeptide |
| 5 | PWWP2A | PWWP domain containing 2A |
| 5 | RANBP17 | RAN binding protein 17 |
| 5 | RASA1 | Ras GTPase-activating protein 1 (GAP) (GTPase-activating protein) (RasGAP) (Ras p21 protein activator) (p120GAP) |
| 5 | ARHGAP26 | Rho GTPase-activating protein 26 (GTPase regulator associated with focal adhesion kinase) (Oligophrenin-1-like protein) (Rho-type GTPase-activating protein 26) |
| 5 | PTTG1 | Securin (Esp1-associated protein) (Pituitary tumor-transforming gene 1 protein) (Tumor-transforming protein 1) (hPTTG) |
| 5 | TLX3 | T-cell leukemia, homeobox 3 (HOX11L2) |
| 5 | TERT | telomerase reverse transcriptase |
| 5 | FER | Tyrosine-protein kinase Fer (EC 2.7.10.2) (Feline encephalitis virus-related kinase FER) (Fujinami poultry sarcoma/Feline sarcoma-related protein Fer) (Proto-oncogene c-Fer) (Tyrosine kinase 3) (p94-Fer) |
| 8 | CHCHD7 | coiled-coil-helix-coiled-coil-helix domain containing 7 |
| 8 | CBFA2T1 | core-binding factor, runt domain, alpha subunit 2;translocated to, 1 (ETO) |
| 8 | COX6C | cytochrome c oxidase subunit VIc |
| 8 | EIF3E | eukaryotic translation initiation factor 3, subunit E |
| 8 | FGFR1 | fibroblast growth factor receptor 1 |
| 8 | HEY1 | hairy/enhancer-of-split related with YRPW motif 1 |
| 8 | KAT6A | Histone acetyltransferase KAT6A (EC 2.3.1.48) (MOZ, YBF2/SAS3, SAS2 and TIP60 protein 3) (MYST-3) (Monocytic leukemia zinc finger protein) (Runt-related transcription factor-binding protein 2) (Zinc finger protein 220) |
| 8 | HOOK3 | hook homolog 3 |
| 8 | IKBKB | inhibitor of kappa light polypeptide gene enhancer in B-cells, kinase beta |
| 8 | NRG1 | neuregulin 1 |
| 8 | NDRG1 | N-myc downstream regulated 1 |
| 8 | NCOA2 | nuclear receptor coactivator 2 (TIF2) |
| 8 | PCM1 | pericentriolar material 1 (PTC4) |
| 8 | PLAG1 | pleiomorphic adenoma gene 1 |
| 8 | RUNX1T1 | Protein CBFA2T1 (Cyclin-D-related protein) (Eight twenty one protein) (Protein ETO) (Protein MTG8) (Zinc finger MYND domain-containing protein 2) |
| 8 | MOS | Proto-oncogene serine/threonine-protein kinase mos (EC 2.7.11.1) (Oocyte maturation factor mos) (Proto-oncogene c-Mos) |
| 8 | RSPO2 | R-spondin 2 |
| 8 | RUNXBP2 | runt-related transcription factor binding protein 2 (MOZ/ZNF220) |
| 8 | TCEA1 | transcription elongation factor A (SII), 1 |
| 8 | MAFA | Transcription factor MafA (Pancreatic beta-cell-specific transcriptional activator) (Transcription factor RIPE3b1) (V-maf musculoaponeurotic fibrosarcoma oncogene homolog A) |
| 8 | LYN | Tyrosine-protein kinase Lyn (EC 2.7.10.2) (Lck/Yes-related novel protein tyrosine kinase) (V-yes-1 Yamaguchi sarcoma viral related oncogene homolog) (p53Lyn) (p56Lyn) |
| 8 | MYC | v-myc myelocytomatosis viral oncogene homolog (avian) |
| 8 | WISP1 | WNT1-inducible-signaling pathway protein 1 (WISP-1) (CCN family member 4) (Wnt-1-induced secreted protein) |
| 8 | WHSC1L1 | Wolf-Hirschhorn syndrome candidate 1-like 1 (NSD3) |

† Data are derived from the Sanger Cancer Genome Census and the Uniprot list of proto-oncogenes.

